# Haplotype editing with CRISPR/Cas9 as a therapeutic approach for dominant-negative missense mutations in *NEFL*

**DOI:** 10.1101/2024.12.20.629813

**Authors:** Poorvi H. Dua, Bazilco M. J. Simon, Chiara B.E. Marley, Carissa M. Feliciano, Hannah L. Watry, Dylan Steury, Abin Abraham, Erin N. Gilbertson, Grace D. Ramey, John A. Capra, Bruce R. Conklin, Luke M. Judge

## Abstract

Inactivation of disease alleles by allele-specific editing is a promising approach to treat dominant-negative genetic disorders, provided the causative gene is haplo-sufficient. We previously edited a dominant *NEFL* missense mutation with inactivating frameshifts and rescued disease-relevant phenotypes in induced pluripotent stem cell (iPSC)-derived motor neurons. However, a multitude of different *NEFL* missense mutations cause disease. Here, we addressed this challenge by targeting common single-nucleotide polymorphisms in cis with *NEFL* disease mutations for gene excision. We validated this haplotype editing approach for two different missense mutations and demonstrated its therapeutic potential in iPSC-motor neurons. Surprisingly, our analysis revealed that gene inversion, a frequent byproduct of excision editing, failed to reliably disrupt mutant allele expression. We deployed alternative strategies and novel molecular assays to increase therapeutic editing outcomes while maintaining specificity for the mutant allele. Finally, population genetics analysis demonstrated the power of haplotype editing to enable therapeutic development for the greatest number of patients. Our data serve as an important case study for many dominant genetic disorders amenable to this approach.

## INTRODUCTION

The discovery and advancement of gene-editing technologies are revolutionizing how scientists and clinicians approach treatments for genetic diseases^1^. The CRISPR-Cas system is a powerful gene-editing tool with two fundamental components, a programmable guide RNA (gRNA) that encodes sequence complementary to a genomic locus, and a Cas enzyme that induces a double-stranded break (DSB) in DNA when its associated gRNA binds to the target site^2^. DSB repair pathways can generate short insertions and deletions (indels) at the cut site that disrupt the DNA sequence; this simplest form of gene editing is the basis of the first approved gene editing therapy for hemoglobinopathies^2^. Hemoglobinopathies and other conditions tested in ongoing clinical trials are representative of the small number of disorders where the disruption of both alleles of a coding gene or regulatory element could lead to therapeutic benefit with minimal toxicity (NCT04601051, NCT06128629, NCT05120830). However, the majority of genetic disorders will require more complex and precise editing strategies. For example, autosomal dominant missense mutations, which are highly prevalent in neurologic, ophthalmologic, and cardiovascular diseases, require allele-specific precision because disrupting both the mutant and normal allele would impair cellular function. We and other groups have demonstrated that allele-specific editing to selectively disrupt a mutant allele is a promising strategy for dominant-negative or gain-of-function disorders where a single allele is adequate for normal function^3–6^.

An example of an autosomal dominant disorder amenable to allele-specific editing is Charcot-Marie-Tooth disease type 2E (CMT2E), a genetic neuropathy with no current treatment. CMT2E is caused by mutations in the *NEFL* gene^7^, which encodes the neurofilament light chain protein (NF-L) that forms the core of hetero-oligomeric neurofilaments critical for axonal growth and function^8^. Missense mutations in *NEFL* act via dominant-negative mechanisms whereby mutant NF-L interferes with the proper assembly, transport, or function of neurofilament oligomers^9–12^. Interestingly, loss-of-function mutations in *NEFL* are recessive, and people heterozygous for these mutations exhibit normal neurological function, evidence that selective inactivation of dominant missense mutations in *NEFL* should be well tolerated in patients^13,14^.

Motor neurons differentiated from patient-derived induced pluripotent stem cells (iPSC) have been demonstrated to recapitulate CMT2E pathology in vitro^15,16^. We previously utilized this model to demonstrate rescue of disease phenotypes after inactivating the *NEFL* N98S mutant allele via selective introduction of frameshift-producing indels at the mutation site^4^. However, patients with CMT2E can present with any one of more than 50 different causative missense mutations distributed over all *NEFL* exons (Figure 1a). This diversity of causative mutations motivated our search for a mutation-agnostic editing strategy that would be therapeutic for a greater proportion of the patient population.

**Figure 1:**
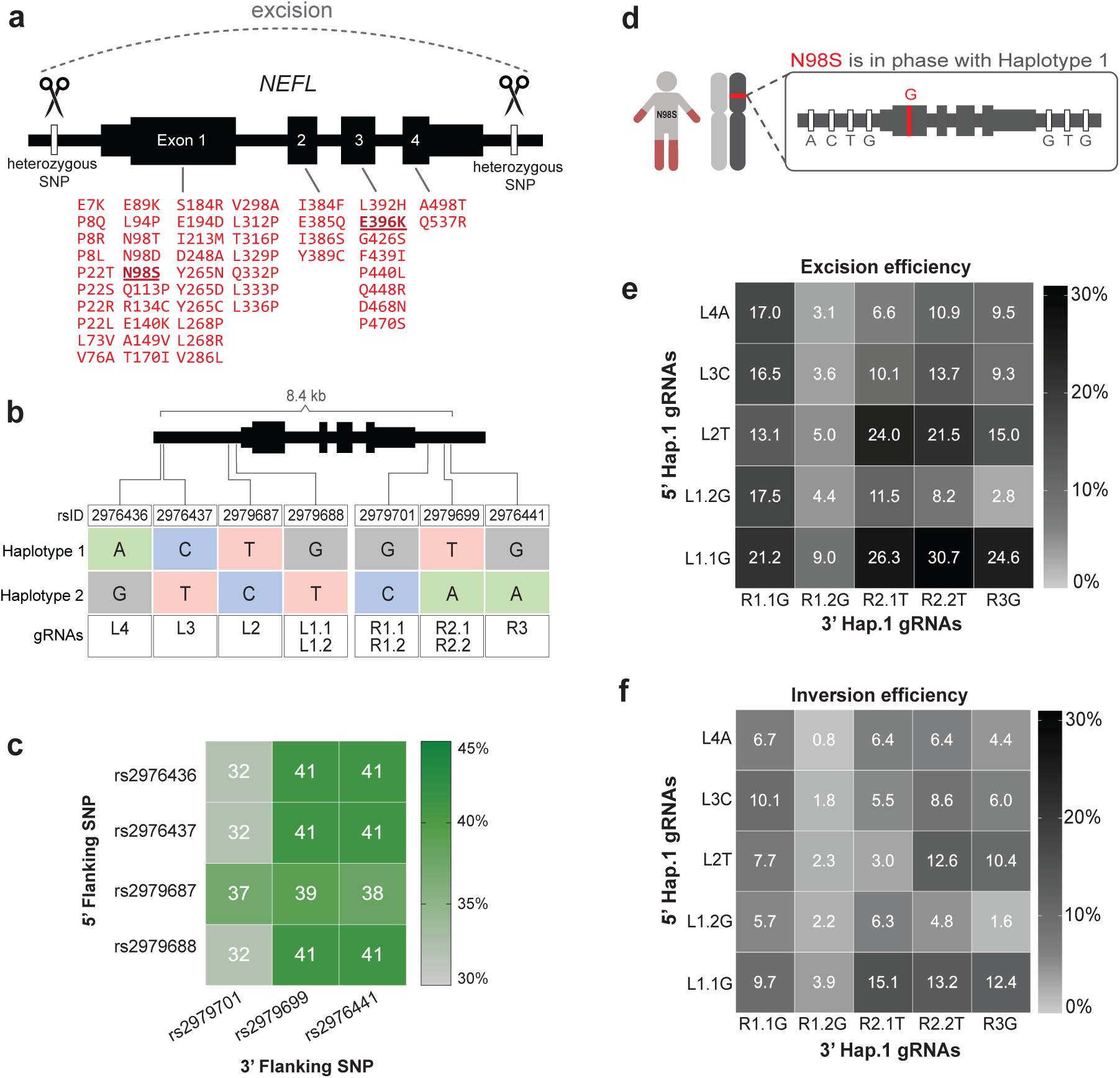
Common SNPs flanking the *NEFL* locus can be targeted to excise the N98S dominant allele. **(a)** Schematic of mutation-agnostic approach to excise any pathogenic *NEFL* mutation (indicated in red) using pairs of SNP-targeting gRNA- HiFiCas9 (scissors). **(b)** Panel of gRNAs targeting SNPs immediately 5’ (L) or 3’ (R) of *NEFL* and numbered by increasing distance from the gene body. Decimal numbers distinguish gRNA targeting the same SNP using different protospacer-adjacent motifs (PAM). Variant sequences shown define two haplotypes present in each heterozygous cell line. **(c)** Heatmap showing the percentage of individuals from 1kGP heterozygous for pairs of SNPs targeted by our panel of gRNAs. **(d)** Schematic of the mutation in N98S-P2 in phase with SNPs defining haplotype 1. **(e-f)** Heatmaps of excision (e) and inversion (f) frequencies measured by ddPCR after paired gRNA-HiFiCas9 transfections in N98S-P2 iPSCs. Data represent the mean of duplicate transfections.

In this study, we achieved mutation-agnostic and allele-specific editing via a dual-gRNA CRISPR system that targets commonly inherited non-coding variants in the human population. CRISPR editing using paired gRNA can efficiently induce large-scale excision or inversion of the intervening sequence^17–20^. Excision-based editing has progressed to clinical trials to revert inherited retinal dystrophy or eliminate HIV proviral genomes, highlighting the therapeutic potential of this concept (NCT05144386, NCT03872479). Furthermore, allele-specific excision has been proposed as a therapeutic strategy for repeat expansion disorders like FTD-ALS and Huntington’s Disease^21–23^ and dominant missense mutations causing corneal dystrophy^24,25^.

Here, we evaluate allele-specific editing in multiple patient-derived iPSC cell lines with novel molecular assays and clonal analysis of editing outcomes to rigorously interrogate haplotype editing in a robust disease model. Our results provide proof-of-principle for haplotype editing as a therapeutic strategy for any dominant missense mutation causing CMT2E. While our studies are focused on mutations in *NEFL* and their effects on motor neurons, the approach and discoveries we describe are broadly applicable to a wide variety of dominant genetic disorders.

## RESULTS

### Mutation-specific gRNAs induce indels with varying efficiency

Our previous study developed a robust iPSC-derived model of genetic motor neuropathy in which we established allele-specific editing as a promising therapeutic strategy for dominant mutations in *NEFL*^4^. That study used a single CMT2E iPSC line with the N98S missense mutation (N98S-P1). To test whether the same approach would generalize to other *NEFL* mutations and genetic backgrounds, we generated CMT2E iPSC lines from an unrelated N98S patient of different sex and ethnic background (N98S-P2), and from two related patients with the E396K missense mutation (E396K-P1, E396K-P2). We engineered these patient-derived iPSCs with integrated, inducible, and isogenic (i^3^) transcription factors so we could rapidly and efficiently differentiate them into lower motor neurons (i^3^LMNs)^26^.

Our established allele-specific editing strategy used a single mutation-specific gRNA and high-fidelity Cas9 derived from S. pyogenes (HiFiCas9)^27^ that allowed for the introduction of frameshift indels in the coding region of the mutant *NEFL* allele, which in turn induced nonsense-mediated decay of the mutant transcript. This gRNA-HiFiCas9 ribonucleoprotein (RNP) induced similarly efficient editing of the mutant allele in the N98S-P2 iPSCs (Figure S1a, S3a-c). We isolated a clonal iPSC line with a frameshift indel at the N98S mutation site (N98S- P2-fs) and separately generated an isogenic corrected line in the N98S-P2 background as a control (N98S-P2-cor) as described previously (Figure S1a-c)^4^. When we differentiated the frameshift line into i^3^LMNs, we saw that the expression of total *NEFL* mRNA was reduced by approximately 50% compared to unedited N98S-P2 control, with a near-total extinction of the mutant allele (Figure S2a-b). Our group previously developed an automated imaging analysis assay to measure the pathologic NF-L accumulation in the cell bodies of N98S mutant motor neurons^4^. Using this assay, we confirmed that the allele-specific frameshift rescued the pathologic phenotype, consistent with our original results in the N98S-P1 patient line (Figure S2c). Additionally, we confirmed that spontaneous action potentials were similar in clonal N98S- P2-fs i^3^LMNs compared to unedited N98S-P2 and N98S-P2-corrected i^3^LMNs (Figure S2d).

We attempted to extend this editing strategy to the E396K mutation (Figure S3a). However, we detected only minimal indels at the target locus in both E396K patient lines, as measured by amplicon sequencing (Figure S3b-c). We confirmed the nuclease activity of the E396K gRNA-HiFiCas9 RNP using an in vitro cleavage assay (Figure S3d). These findings suggest that double-strand breaks at the E396K site may be prone to error-free repair in iPSCs.

Considering that patients with CMT2E have been reported with any one of > 50 different missense mutations in *NEFL* (Figure 1a), and that new mutations continue to be discovered, we reasoned that optimizing gRNAs for each possible mutation would be a laborious and impractical approach. We therefore looked for an alternative approach that would be allele-specific, but also mutation-agnostic.

### Haplotype-specific gRNA pairs excise the *NEFL-*N98S allele with varying efficiency

To find other ways to inactivate *NEFL* mutant alleles, we examined whole genome sequencing of our four patient cell lines. We uncovered many heterozygous variants flanking *NEFL* in 3 out of 4 of our cell lines (Figure S4). Since these variants are in non-coding intergenic regions, indels at any one of the variant sites alone would be unlikely to inactivate the allele.

Therefore, we designed gRNAs to target pairs of variants flanking *NEFL* with the goal of excising the entire mutant gene (Figure 1b). We validated the relevance of these allele-specific gRNAs for the broader human population by analyzing phased genomes from the 1000 Genomes Project (1kGP)^28^. We found that 32-41% of individuals are heterozygous for pairs of common single nucleotide polymorphisms (SNP) that could be targeted for allele-specific gene excision (Figure 1c). Finally, we determined that the disease mutation in N98S-P2 is in phase with the reference allele for each variant, which we refer to as haplotype 1 (Figure 1d). The mutation in E396K-P1 and E396K-P2 is in phase with the alternate alleles, which we refer to as haplotype 2 (Figure S5a).

We tested the haplotype excision strategy first with N98S-P2 so we could compare it with our original N98S mutation-targeting strategy. We paired each gRNA upstream of *NEFL* with each downstream gRNA and transfected N98S-P2 iPSCs with the resulting SNP-specific HiFiCas9 RNPs. We measured the frequency of excision by ddPCR using our established methodology^17^ and found that it varied from 2.8% to 30.7% depending on the gRNA pair (Figure 1e). We also detected measurable levels of inversion in the same samples, ranging from 0.8% to 15.1% (Figure 1f). The most efficient combinations produced combined excision and inversion editing at >40% of total alleles, with 50% being the maximum expected if editing is perfectly specific for the target allele. Pairs containing the L1.1G gRNA induced high levels of both excision and inversion, whereas pairs containing the R1.2G gRNA induced remarkably low levels of both. Overall, these findings show that efficient haplotype-specific editing with gRNA- HiFiCas9 pairs is feasible at the *NEFL* locus. Next, we sought to investigate the therapeutic potential of this strategy in our i^3^LMN disease model of CMT2E.

### Haplotype-specific excision of N98S and E396K alleles restores NF-L subcellular distribution in patient iPSC-derived motor neurons

To examine the effect of allele-specific haplotype editing on *NEFL* mRNA expression, we focused on gRNA targeting SNPs closest to the 5’ and 3’ ends of *NEFL* (L1.1 and R1.1). This pair was one of the highest in editing efficiency, while also producing the smallest possible excision. We transfected N98S-P2 iPSCs with L1.1G and R1.1G RNPs targeting haplotype 1 (Figure 2a) and differentiated them into i^3^LMNs. We measured mRNA transcript levels by quantitative RT-ddPCR and found that the edited population had decreased total *NEFL* mRNA compared to the unedited control, with an expected decrease in the fraction of mutant mRNA (Figure 2b-c). To determine the effect of excision vs. inversion on *NEFL* expression, we isolated clonal iPSC lines with excision or inversion of the N98S allele and differentiated them into i^3^LMNs. As expected, neurons derived from the excision clone showed no expression of the N98S allele and an overall 50% decrease in total *NEFL* mRNA (Figure 2b-c). Interestingly, neurons derived from the inversion clone demonstrated minimal change in total or mutant *NEFL* mRNA, suggesting that inversion induced by this CRISPR editing strategy fails to separate the gene’s coding region from its promoter and other regulatory elements.

**Figure 2:**
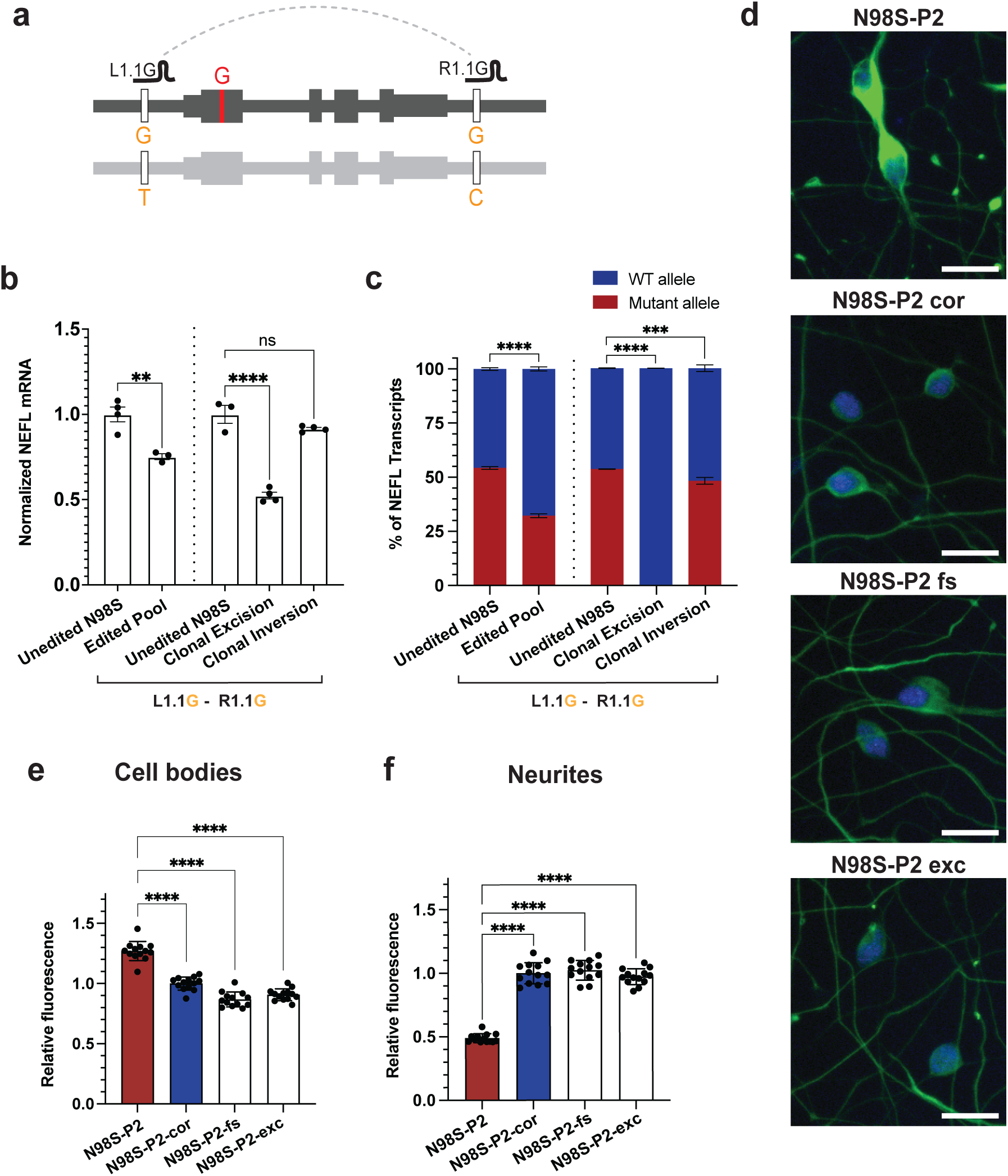
Allele-specific excision of *NEFL* inactivates mutant allele expression and rescues phenotypes in patient iPSC-motor neurons. (**a**) Schematic of SNP-targeted excision of N98S allele. Edited iPSCs from patient N98S-P2 were differentiated into i^3^LMNs, RNA isolated on Day 7 and measured by quantitative RT-ddPCR. **(b)** Total *NEFL* expression relative to *GAPDH* and normalized to unedited control. **(c)** Relative allelic expression as determined by ddPCR using a heterozygous SNP in the 3’ UTR (rs2976439). Bar graphs for (b) and (c) indicate mean +/- S.E.M. of biological replicates normalized to unedited control. **(d)** Representative images of day-7 i^3^LMNs stained with anti-NF-L (green) and anti-HB9 (blue) antibodies. Scale bars = 20 µM. **(e)** Quantification of mean NF-L fluorescence intensity in cell bodies. **(f)** Quantification of total NF-L fluorescence intensity in neurites normalized to β3-tubulin positive neurite area. Bar graphs for (e-f) represent mean +/- S.E.M. of biological replicates per cell line normalized to isogenic control (N98S-P2-cor). Each data point represents mean values from 15 images per biological replicate. ** p < 0.01, *** p < 0.001, **** p < 0.0001, ns p > 0.05.

To confirm that haplotype editing is also effective at targeting haplotype 2, we repeated the excision in E396K-P2 iPSCs (Figure S5b), using the L1.1T and R1.1C gRNAs, which are specific to haplotype 2. We differentiated the population of edited E396K-P2 iPSCs into i^3^LMNs and again observed decreased total *NEFL* mRNA compared to the unedited control, with a corresponding decrease in the fraction of mutant mRNA (Figure S5c). When examining clones of edited cells, we confirmed our previous finding that the excision abolished transcription of the mutant E396K allele and reduced total *NEFL* mRNA by 50%, while the inversion produced minimal changes in mutant or total *NEFL* expression (Figure S5c-d).

One manifestation of *NEFL*-associated pathology is the accumulation of neurofilaments in the cell body of motor neurons^4,15,29^. To determine whether allelic inactivation via excision rescued this phenotype, we stained i^3^LMNs derived from N98S-P2 iPSCs for NF-L and the nuclear motor neuron marker HB9, and measured the NF-L fluorescence signal with our established image analysis pipeline^4^. We compared N98S-P2-excision to the isogenic frameshift, corrected, and unedited controls, and found that excision was as effective as frameshift at normalizing NF-L intensity in motor neuron cell bodies (Figure 2d, S6a). We performed the same analysis comparing E396K-P2-excision with E396K-P2-inversion, unedited control, and an unrelated healthy control line (WTC). We observed a similar pattern as in the N98S series, although the E396K phenotype was more subtle (Figure S6b-c) consistent with functional comparisons of the effects of these mutations on neurofilament assembly^12^.

In N98S mutant mice, aggregation of neurofilaments in neuronal cell bodies is associated with severe depletion of neurofilaments in axons^30^. We questioned whether we could detect a similar neurofilament protein depletion in neurites of our motor neurons, and if inactivation of the mutant allele could restore this distribution despite the decrease in total transcript levels. To test this hypothesis, we modified our staining protocol to simultaneously measure NF-L in cell bodies and neurites from a single imaging assay by including anti-β3 tubulin to identify neurites and replacing anti-HB9 with a general nuclear stain to identify cell bodies (Figures S7, S8).

Furthermore, we incorporated a fully automated microscopy platform to increase the throughput and content of our dataset in an unbiased manner. We designed complementary CellProfiler^31^ pipelines to quantify fluorescence intensity in the cell bodies and neurites from this imaging platform (Figures S7, S8). This analysis recapitulated our finding that frameshift and excision of the N98S allele similarly rescue the pathological accumulation of NF-L in the cell body, confirming the validity of this assay (Figure 2e). Consistent with the in vivo results from mice, we observed a dramatic depletion of NF-L in N98S neurites compared to the isogenic corrected control (Figure 2f). Furthermore, excision or frameshift inactivation of N98S restored NF-L levels within neurites (Figure 2f). We previously showed that inactivation of *NEFL*-N98S leads to reduced total NF-L protein in i^3^LMNs^4^. Our findings here demonstrate that the reduced accumulation of NF-L in cell bodies also reflects redistribution to other cellular locations, with likely important functional consequences.

To evaluate whether off-target genome editing could confound the biological behavior of edited clones, we sequenced bioinformatically predicted off-target sites in each clonal line derived from N98S-P2 and E396K-P2 and found no evidence of CRISPR-induced mutations (Table S1). Additionally, we determined that the excision or inversion of *NEFL* did not affect the expression of *NEFM*, which encodes the co-regulated neurofilament-medium chain that is adjacent to the *NEFL* locus (Figure S9). Finally, we performed electrophysiological analysis on differentiated motor neurons using a multi-electrode array. Spontaneous action potentials were similar in all excision and inversion clones and their unedited N98S-P2 and E396K-P2 controls, suggesting that editing did not disrupt neuronal electrophysiological function (Figure S10).

### Including a single-stranded bridging oligonucleotide boosts excision efficiency by 30–50%

Although haplotype editing at our target SNP sites induced significant levels of both excisions and inversions, only the excision inactivated the mutant allele. Therefore, we investigated methods to increase the frequency of excision outcomes. Previous work indicated that a single-stranded oligonucleotide donor (ssODN) could induce long-range excision in the context of a single DSB induced by zinc-finger nucleases^32^. Reasoning that a similar strategy might also enhance excisions produced by paired Cas9 RNPs, we transfected N98S-P2 and E396K-P2 iPSCs with L1.1 and R1.1 RNP and included a 60-nucleotide ssODN designed to mimic the excision repair outcome. We measured both excision and inversion frequency by ddPCR and found a near 50% increase in excision frequency in N98S-P2 iPSC and a near 33% increase in E396K-P2 iPSC while inversion frequency showed no significant change in either patient line (Figure 3). We also investigated indel outcomes at the individual target sites, and found they varied inconsistently with addition of the bridging oligo (Figure S11a).

**Figure 3:**
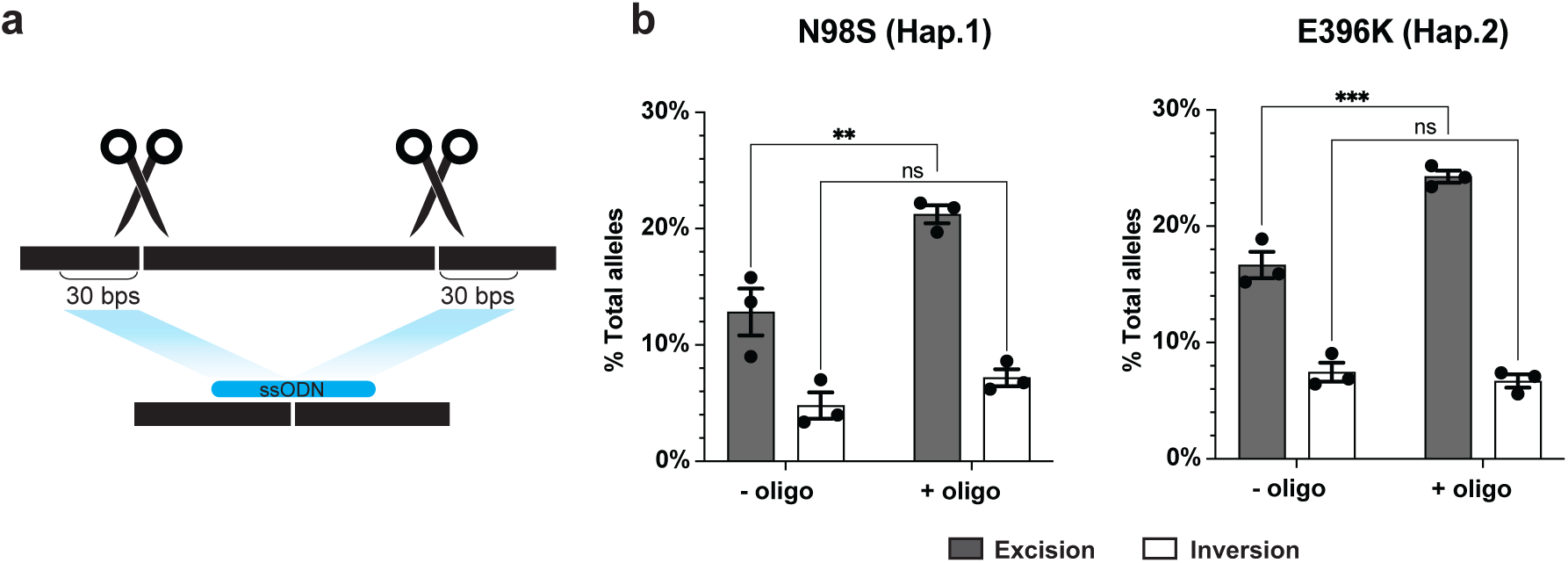
Excision frequency is increased by single-stranded oligonucleotide (ssODN) bridging donors. **(a)** Schematic of bridging ssODN design complementary to the expected excision repair product. **(b)** Excision and inversion frequencies +/- ssODN quantified by ddPCR after transfection of L1.1-R1.1 RNPs in N98S-P2 and E396K-P2 iPSC. Bar graphs represent the mean +/- S.E.M. of replicate transfections. ** p < 0.01, *** p < 0.001, ns p > 0.05.

To confirm that the effect of the bridging oligo is not unique to this specific excision outcome, we repeated the experiment with a smaller excision removing only the first exon. We observed a similar increase in excision efficiency and no effect on inversion (Figure S11b). We decreased the length of the ssODN to 30 nucleotides, which eradicated the boost in efficiency, while increasing the length of the ssODN to 120 nucleotides or adding stabilizing chemical modifications did not significantly improve the frequency of excisions (Figure S11c-d). These results indicate that the simple addition of a 60-nucleotide ssODN can increase excision efficiency without suppressing inversions.

### Incorporating a biallelic intronic gRNA enables both excisions and inversions to inactivate the mutant allele

We hypothesized that mutant *NEFL* expression was unchanged after L1.1-R1.1 inversion because the transcription start site remains linked with its core promoter. To increase the efficiency of our editing approach, we sought to design gRNA pairs that would cause both excisions and inversions to disrupt transcription of the mutant alleles. We reasoned that we could achieve this goal by combining either one of our flanking gRNAs (L1.1 or R1.1) with an intragenic gRNA (Figure 4a). However, intragenic SNPs are less common in the population and our cell lines, limiting our ability to design allele-specific intragenic gRNAs. Thus, we designed gRNAs common to both alleles (biallelic) in the first intron of *NEFL* and paired them with our SNP-specific gRNAs for partial gene excision and inversion (Figure 4a). Since biallelic gRNAs are expected to induce indels on the wild-type allele, we first assessed whether indels in the targeted region of the intron would affect *NEFL* expression. We tested two biallelic gRNAs in the first intron that generated indels at up to 89% efficiency (Figure S12a). Differentiation of the edited populations to i^3^LMNs demonstrated maintained *NEFL* expression despite the high frequency of intronic indels (Figure S12b).

**Figure 4:**
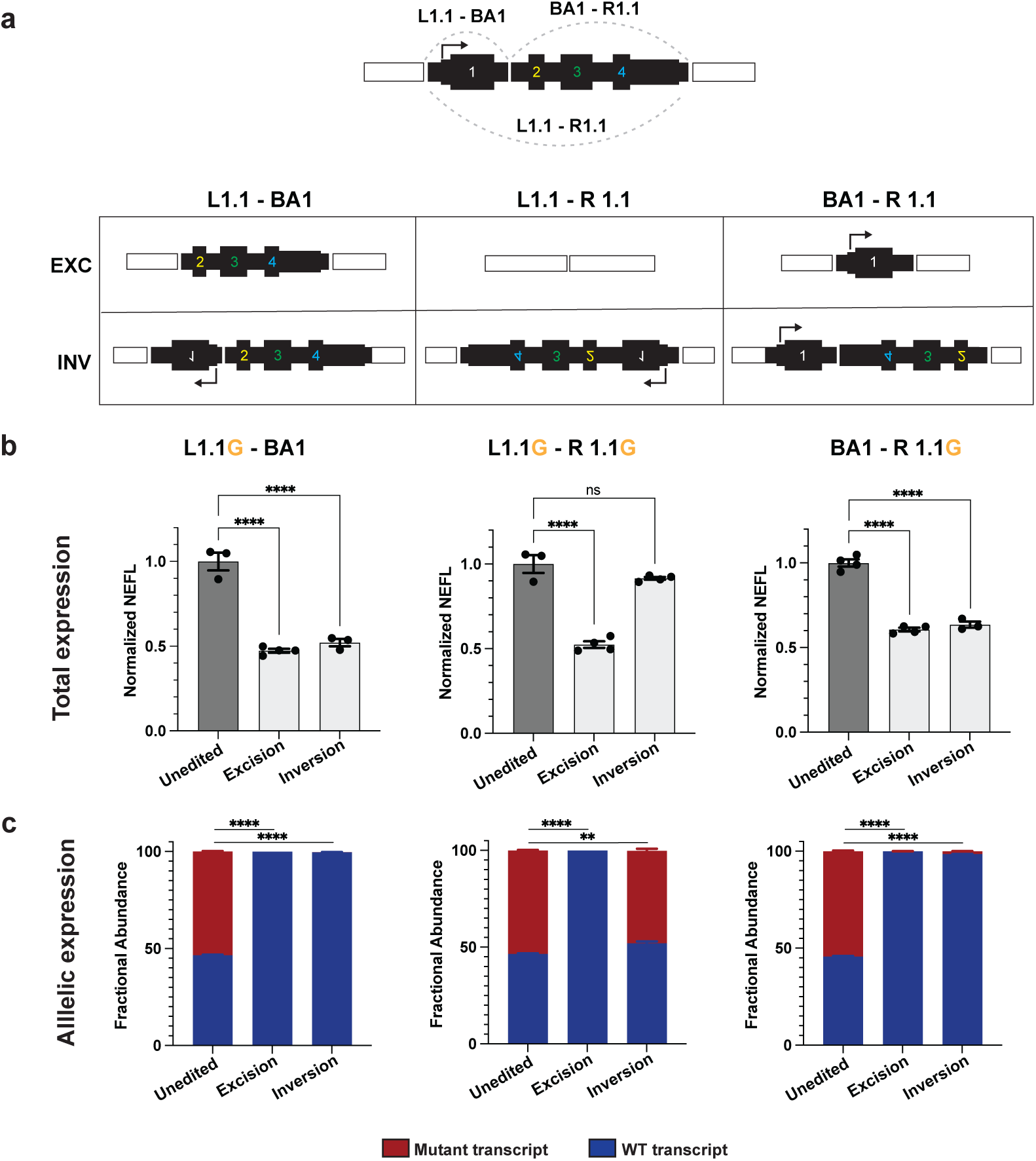
gRNA pairs that include an intronic gRNA induce excisions and inversions that both inactivate mutant alleles. **(a)** Schematic of predicted editing outcomes with various combinations of L1.1, R1.1 and intronic biallelic (BA1) gRNAs. N98S-P2 was transfected with each RNP pair and clonal iPSC lines were isolated with each predicted outcome and differentiated into i^3^LMNs. RNA was isolated on Day 7 and measured by quantitative RT-ddPCR. **(b)** Total *NEFL* expression relative to *GAPDH* and normalized to unedited control. **(c)** Relative allelic expression was assessed via allele discrimination ddPCR using a heterozygous SNP in the 3’ UTR (rs2976439). Bar graphs represent the mean +/- S.E.M. of biological replicates. ** p < 0.01, **** p < 0.0001, ns p > 0.05.

Next, we paired the biallelic gRNA with highest indel-inducing efficiency (BA1) with each of the SNP-specific gRNAs we used previously (L1.1 or R1.1) to determine whether partial gene excision and inversion would lead to mutant allele inactivation. In the case of the L1.1-BA1 excision, the transcription start site (TSS) and entire first exon would be deleted, whereas inversion would disrupt the orientation of the TSS and first exon from the rest of the gene. In the case of the BA1-R1.1 excision, the final three exons would be deleted, whereas inversion would disrupt the orientation of the final three exons including the translation and transcription terminator sequences (Figure 4a). We transfected N98S-P2 iPSCs with RNP containing the L1.1G - BA1 gRNA pair and the BA1 - R1.1G gRNA pair, isolated clones with our desired excisions and inversions, and differentiated these clones into i^3^LMNs. With both L1.1-BA1 and BA1-R1.1 conditions, excision led to 50% reduction in total *NEFL* mRNA and no detectable expression of the N98S allele (Figure 4b-c). More notably, inversion also led to a 50% reduction in total *NEFL* transcripts and no detectable expression of the mutant allele. We applied this strategy to the E396K line and obtained comparable results (Figure S13).

### High-efficiency and -specificity therapeutic editing can be achieved either with a pair of allele-specific gRNAs or with a biallelic gRNA paired with an allele-specific gRNA

Our ultimate goal is to optimize an allele-specific editing strategy that produces the most therapeutic events (i.e. inactivation of the mutant *NEFL* allele) with the lowest risk of editing the normal allele. To this end, we first compared the efficiency and specificity of edits generated with L1.1-BA1, L1.1-R1.1, and BA1-R1.1. We transfected each of these three gRNA pairs into N98S-P2 and E396K-P2 iPSCs along with a 60 nt bridging ssODN (following our previous design) and measured the excision and inversion efficiency by ddPCR in polyclonal iPSC populations (Figure 5a). In both N98S-P2 and E396K-P2 iPSCs, the pairs that included the biallelic gRNA produced more combined excisions plus inversions than did the pair of allele-specific gRNAs (Figure 5b). Notably, the gRNA pair L1.1G-BA1 in N98S-P2 produced combined excision and inversion edits in approximately 60% of alleles. Since allele-specific editing should ideally occur on a maximum of 50% of total alleles, we hypothesized that the introduction of a biallelic gRNA was leading to undesired excision and/or inversion of the non-target allele. Thus, we sought to thoroughly investigate the specificity of each gRNA combination.

**Figure 5:**
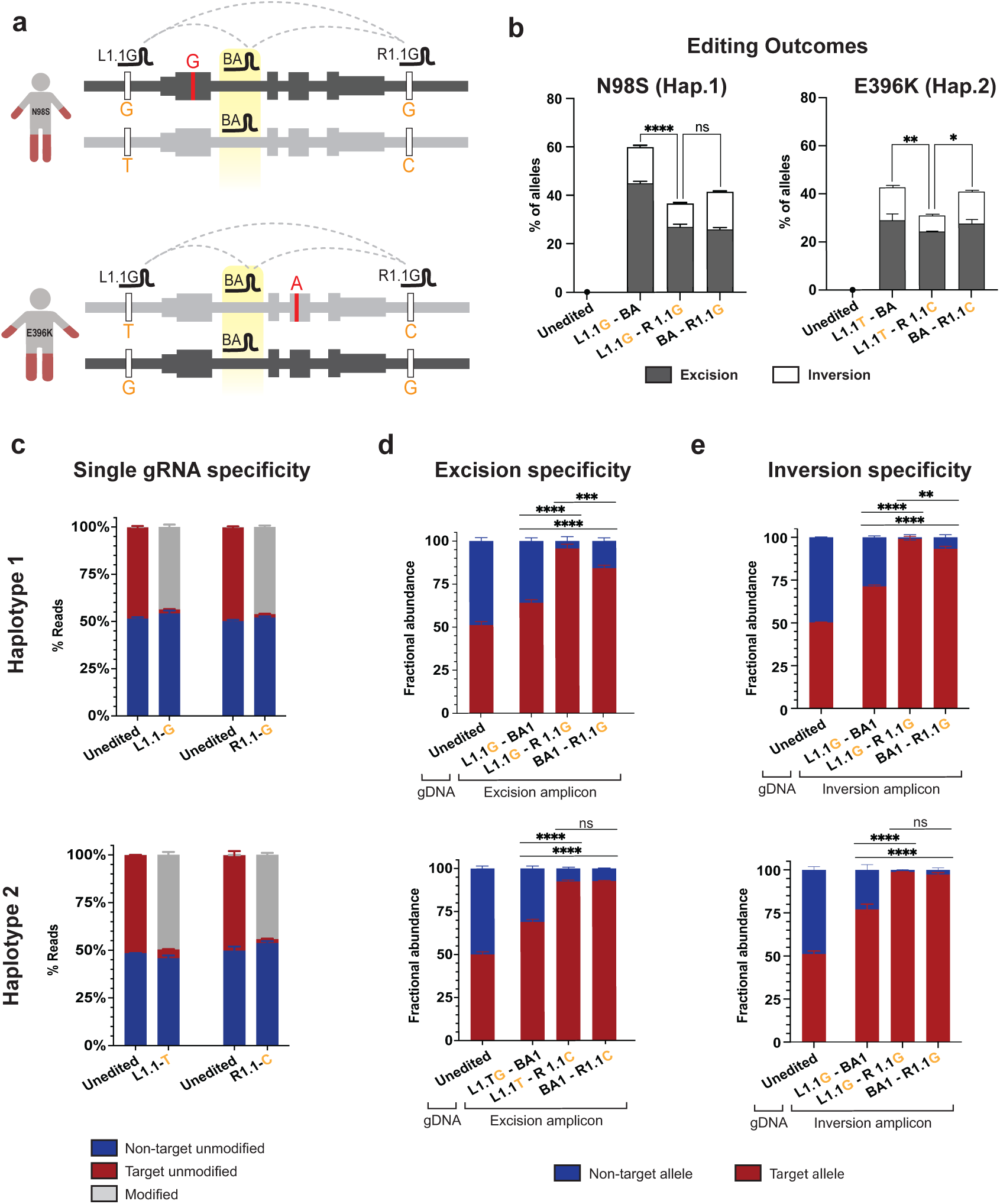
Optimization of allele-specific haplotype editing. **(a)** Schematic of haplotype-specific editing for N98S-P2 (haplotype 1) and E396K-P2 (haplotype 2) with various combinations of L1.1, R1.1 and intronic biallelic (BA1) gRNAs. **(b)** Quantification of excision and inversion frequency via ddPCR. **(c)** Quantification of allele-specific editing with individual gRNAs via NGS amplicon sequencing. **(d-e)** Quantification of allele specificity of excision and inversion via a ddPCR allele discrimination assay to a linked heterozygous SNP (rs2979685) located 5’ of the *NEFL* gene. Unedited controls demonstrate equal abundance of variant alleles in gDNA at baseline while edited samples represent fractional abundance of alleles in excision or inversion events. All graphs represent mean +/- S.E.M. of replicate transfections. * p < 0.05, ** p < 0.01, *** p < 0.001, **** p < 0.0001, ns p > 0.05.

First, we evaluated the individual editing efficiencies and specificities of eight SNP-specific gRNAs: the gRNA set that targeted haplotype variants of L1.1 and R1.1, as well as another set targeting the same two SNPs but with different PAM and protospacer sequences (L1.2 and R1.2, Figure S14a). We transfected N98S-P2 or E396K-P2 iPSCs with RNP targeting their mutant haplotype and measured indels via deep amplicon sequencing, followed by analysis using Crispresso2. This program includes a quantification feature that deconvolutes sequencing reads by allele based on the sequence of the heterozygous variant harbored by each allele^33^. Both haplotype versions of L1.1 and R1.1 were efficient at producing indels, with no evidence of editing at the non-target allele (Figures 5c, S14b-c). Interestingly, both haplotype versions of L1.2 induced significant editing of the non-target allele despite the SNP being located in the PAM-proximal seed region known to be important for single nucleotide specificity (Figure S14a- b)^34^. Both haplotype versions of R1.2 induced fewer indels than R1.1 (Figure S14b). This low indel efficiency is consistent with the low excision and inversion frequencies we observed using R1.2G in the initial arrayed screen (Figure 1e-f).

Since L1.1 and R1.1 gRNAs were highly efficient and specific when tested individually, we tested their specificity when paired to produce excisions and inversions. To rigorously measure specificity, we developed new variations of our ddPCR-based methodology. We first designed PCR assays to amplify each excision and inversion breakpoint along with a flanking SNP; the amplified fragments were isolated by gel electrophoresis and used as the template for allele discrimination ddPCR (Figure S15). As expected, excision and inversion with the combination of two allele-specific gRNA were highly specific for the target haplotype in both cell lines, with 92-99% of edits occurring on the target allele (Figure 5d-e). On the other hand, pairing L1.1 with BA1 resulted in a significant decrease in the specificity of either excision or inversion to 64-77%. This result is consistent with the higher overall editing efficiency we observed with this pair (Figure 5b). Interestingly, the specificity of excision and inversion was relatively preserved when pairing R1.1 with BA1 (84-97%). In particular, targeting haplotype 2 in the E396K-P2 cell line with BA1-R1.1C showed equivalent specificity to the allele-specific pair L1.1T-R1.1C (Figure 5d-e), indicating that a single SNP-specific gRNA paired with a biallelic gRNA can be sufficient for highly specific allelic excision and inversion in some cases. Importantly, the differential specificity of L1.1 and R1.1 for excision and inversion when combined with BA1 was not predicted by testing the specificity of the individual gRNAs and was only revealed by our novel assay.

To confirm these results with a separate complementary assay, we designed a multiplexed digital PCR assay with four differentially labeled probes (Figure S16). This assay recapitulated our previously described method utilizing the linkage of an allele-discrimination assay to an excision-specific amplicon to measure allele-specificity in excisions^17^, but with greater precision due to the use of non-overlapping fluorophores. Furthermore, the multiplexed digital PCR allowed us to simultaneously quantify efficiency and specificity (Figure S16, S17). The results corroborated data from the previous separate assays for efficiency and specificity in a single-step reaction without gel electrophoresis. This method processes the complete assay in 96- well format in 2 hours, enabling high-throughput arrayed screens to optimize haplotype editing reagents for both efficiency and specificity.

### Leveraging common haplotypes and paired gRNA editing considerably reduces the number of gRNA-Cas combinations needed to treat the majority of CMT2E patients

To quantify the power of haplotype editing for *NEFL,* we used data from the 1000 Genomes Project^28^ and calculated the percentage of CMT2E patients that could be treated with a given number of unique allele-specific therapies. For mutation-targeted editing we assumed that there are 51 mutations (Figure 1a) that occur with similar frequency in the patient population.

For haplotype editing, we assumed that SNP are distributed similarly in CMT2E patients as in the general human population and calculated the percentage of individuals heterozygous for individual SNP or pairs of SNP flanking the entire gene (Figure S18, S19). Finally, assuming that each SNP can be targeted with an optimized Cas enzyme and gRNA, we calculated the percentage of individuals who could be treated with increasing numbers of Cas/gRNA (Figure 6). Using mutation-specific gRNAs to induce indels, 26 different Cas/gRNA would be required to treat > 50% of patients, whereas for haplotype editing with pairs of common SNP-targeted gRNA, 8 different Cas/gRNA would achieve a similar yield (Figure 6). The yield of haplotype excision is markedly improved through the incorporation of a constant reagent (like our BA1 gRNA). Pairing BA1 with a single allele-specific Cas/gRNA flanking either side of the gene would allow 9 total Cas/gRNA to treat > 90% of patients.

**Figure 6:**
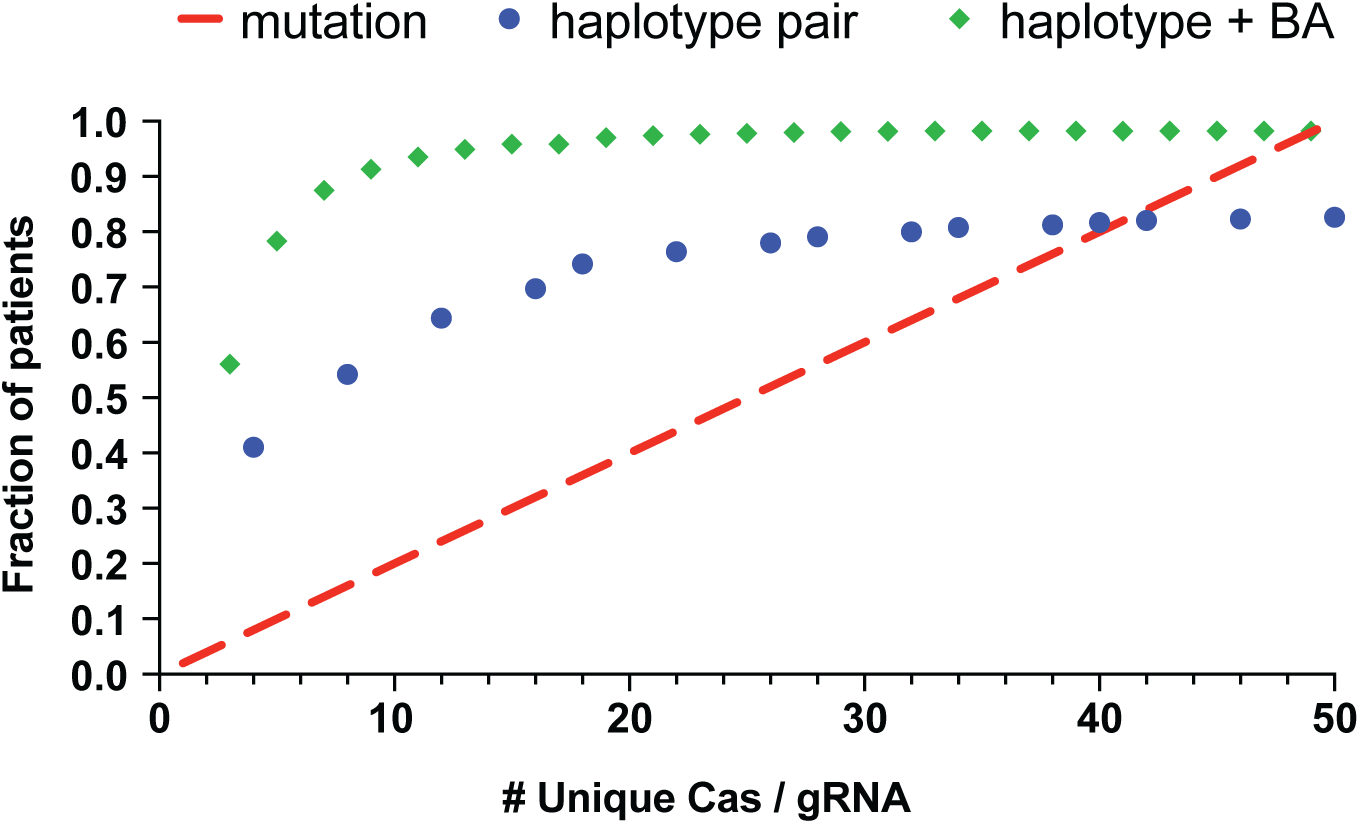
Calculation of patient population coverage with haplotype editing as compared to mutation-targeted editing. Data points illustrate the cumulative number of patients treated with addition of increasing numbers of unique molecular therapies. Red line assumes 51 pathogenic mutations in *NEFL* that are equally prevalent. Blue points represent haplotype excision with two SNP-targeting Cas/gRNAs (some SNPs are present in multiple pairs). Green diamonds represent haplotype excision with one SNP-targeted Cas/gRNA and a constant biallelic Cas/gRNA. Combinations that would excise the neighboring *NEFM* gene are excluded from this analysis.

## DISCUSSION

Genetic disorders caused by autosomal dominant missense mutations represent a significant challenge for gene therapy, which must counteract the disease allele without affecting the normal allele, even though the two may differ by a single nucleotide. We have shown in our previous published work that allele-specific inactivation of the *NEFL* N98S mutation, a dominant missense mutation that causes CMT2E, was achievable with an engineered Cas9 and a mutation-targeting gRNA. This editing strategy rescued CMT2E pathology in patient-derived iPSC-motor neurons, the cell type most affected in the disease. Our current study demonstrated that we can achieve similar success with pairs of guide RNAs targeting common single-nucleotide variants flanking the mutant *NEFL* allele. We showed the validity of this approach for two separate *NEFL* dominant mutations (N98S and E396K) linked with different common haplotypes in the patient population. By taking advantage of naturally occurring and commonly inherited variants, our haplotype editing approach circumvents the need to develop bespoke therapies for each missense mutation, greatly reducing the number of therapies needed to treat the CMT2E patient population. We also anticipate that our approach, along with the improvements we developed to increase its efficacy, will translate to many other diseases that, like CMT2E, are caused by a large number of dominant missense mutations.

The simplest haplotype editing strategy would be to target coding variants to selectively induce frameshift-producing indels on the desired allele, an attractive option for genes with frequent coding variants^35^. However, the majority of coding variants occur at low frequency in the human population, as we observed for *NEFL*^36^. In addition, frameshift editing can fail to achieve the desired outcome for various reasons, including low indel rates that we observed when targeting E396K, or failure to induce nonsense-mediated decay of the transcript with a risk of generating truncated mutant proteins^37^. Excision is a more complex but also widely generalizable strategy to target the abundant non-coding variants in the human genome. Additionally, excision is not limited to coding exons but could be applied to various other sequences, such as repeat expansions and regulatory elements^21,38^.

Other groups have utilized heterozygous variants for allele-specific editing, most of them focused on variants that generate a novel protospacer-adjacent motif (PAM) to recruit the Cas enzyme to the intended allele, rather than relying on mismatches in the gRNA sequence^22–25,35,39^. This “SNP-derived PAM” approach allows specific targeting of the desired allele, but the scope of targetable variants is limited to those that create PAMs and can target only one of the variant alleles. This limits therapeutic impact as many disease-causing missense mutations arise de novo in cis with either ancestral allele. Our strategy incorporated variant-specific sequences in the seed region of the gRNA protospacer to discriminate between alleles, allowing us to target both haplotypes by altering a single nucleotide in the otherwise identical gRNA. This strategy can achieve excellent specificity for both indel and excision editing (Figure 5), although we observed important gRNA-dependent variation. This highlights the importance of our innovations to rigorously evaluate multiple candidates in high throughput, especially as engineered Cas enzymes with minimal PAM requirements allow us to target a larger number of variant sequences^40–42^.

Gene editing therapy for Leber Congenital Amaurosis 10 leverages the combination of excision and inversion for its therapeutic outcomes^18,19^, but many other gene editing studies have ignored the frequency or functional impact of inversions. Here, we showed that inversion is a frequent editing outcome with variable effects on gene expression. We identified multiple strategies to optimize therapeutic editing outcomes, including inversion, while also increasing generalizability for maximum impact on the patient population. Another limitation of previously published studies of allele-specific excision has been the methods for evaluation of excision specificity, and to our knowledge no study has evaluated inversion specificity. Two studies used PCR and Sanger sequencing analysis to provide a qualitative assessment of excision specificity^23,24^. Another used deep amplicon sequencing to evaluate excision specificity by monitoring a heterozygous variant adjacent to the excision site, which provides a high degree of precision and sensitivity^22^. However, due to the limitations of short-read sequencing, only a small number of excisions they tested had variants in requisite proximity for this analysis. Our allele-discrimination digital PCR approach is conceptually similar and highly precise, but more versatile for monitoring haplotype-specific variants distal to the editing site. Furthermore, this novel method allowed us to achieve highly specific excisions and inversions using a single allele-specific gRNA, while revealing differential gRNA specificity that was otherwise undetectable. Our data highlight the utility of pairing allele-specific gRNA with a universal biallelic gRNA targeting any non-coding locus tolerant of small indels. In addition to streamlining the process of therapeutic development (Figure 6), this strategy also creates opportunities to design editing strategies for genes with fewer heterozygous variants.

We demonstrated that a small oligonucleotide donor can increase the frequency of excision editing outcomes. Independent of the therapeutic implications, this simple method will be useful to improve targeted deletions for functional genomics experiments. Going forward, it will be important to investigate these and other strategies for improving excision editing in multiple cell types. For example, our group has already demonstrated that cell-type specific DNA repair pathways lead to differences in indel outcomes in iPSC-derived neurons^43^.

Ultimately, understanding the mechanisms of excision and inversion repair in various disease-relevant cell types will provide insights toward improving the efficiency and precision of therapeutic haplotype editing.

Although *NEFL* is a small and simple gene, there are still thousands of potential SNP combinations to target for haplotype excision (Figure S19). The challenge is even greater for disorders associated with larger and more complex genes, some of which can harbor hundreds of unique disease mutations. We tested a relatively small number of gRNA with a single Cas9 enzyme with promising results, but larger-scale experiments will be important to nominate the best reagents for therapeutic development. Higher throughput experiments testing a wider variety of target loci in multiple genes could also provide insight into genetic and epigenetic factors that maximize the desired editing outcomes. In summary, the innovations described here provide a platform to optimize haplotype editing for a variety of genetic diseases to maximize safety and efficacy for the greatest number of patients.

## METHODS

### Identification of genomic variants and *NEFL* mutation phasing

High molecular weight gDNA was isolated from N98S-P1, N98S-P2, E396K-P1, E396K-P2 iPSC lines using the Blood & Cell Culture DNA Midi Kit (Qiagen 13343). DNA from each line was sent to UC Berkeley QB3 Genomics for whole genome sequencing. Variant calling analysis was completed by Gladstone Institutes Bioinformatics Core resulting in VCF compressed files for each cell line with variants that passed quality check.

Genomic DNA from E396K-P1 and E396K-P2 was extracted using the DNeasy Blood and Tissue Kit (Qiagen 69506). A PCR reaction was performed with primers flanking the E396K mutation and heterozygous variant rs2976439. PCR amplicons were run on a 1% agarose gel and a band of the expected size was extracted using the QIAquick Gel Extraction Kit (Qiagen 28704). The resulting amplicon was cloned into competent bacteria using the TOPO-TA Cloning Kit (Thermo Fisher Scientific, 450641). The transformation reaction was plated on LB-Agar with ampicillin and supplied with 120 µl X-Gal (20 mg/ml) Xgal (Ape Bio A2539) and 40 µl 100 mM IPTG (Gold Bio I2481C). After overnight incubation, white colonies were picked and grown in 5 mL LB Broth supplemented with ampicillin (Fisher BioReagents BP176025). The QIAprep Spin Miniprep Kit (Qiagen 27106) was used to isolate plasmid DNA with amplicon inserts and sent for sequencing. Sequencing of multiple clones confirmed that the E396K mutation was linked to the variant allele of rs2976439.

The phasing of the N98S-P2 line was identified during the characterization of the N98S-P2- frameshift line. A TaqMan SNP genotyping assay targeting the rs79736124 SNP in the 3’ UTR of *NEFL* (ThermoFisher, C_105316276_10) was used to measure allele-specific mRNA expression. We observed a loss of reference allele expression in the clone with N98S-specific frameshift, allowing us to confirm that the N98S mutation in the N98S-P2 patient line is in phase with the reference allele for rs79736124. For both N98S-P2 and E396K-P2, the phasing of more distal SNPs was inferred from population-specific haplotype frequencies using the LDHap tool on the NIH LDLink website (https://ldlink.nih.gov/?tab=ldhap)^44^. Due to strong linkage disequilibrium across the locus (Figure S18), we inferred that the mutation in N98S-P2 was in phase with the reference allele and the mutation in E396K-P2 was in phase with the alternate allele for each variant. All experimental results were consistent with these inferred linkage predictions.

### iPSC reprogramming

Generation of iPSC from peripheral blood mononuclear cells (PBMCs) from the N98S-P2 patient was reported previously^17^. Fibroblasts from the E396K-P1 and E396K-P2 patients were reprogrammed into iPSCs using Stem Cell Technology’s ReproRNA-OKSGM kit (Cat. #05930). Colonies with characteristic iPSC morphology were manually picked and expanded as independent clones sent for karyotypic analysis.

### iPSC maintenance

iPSCs were cultured on Matrigel (Corning, 356231) coated plates at 37 °C, 5% CO2 and 85% humidity. iPSCs were fed mTeSR Plus (Stem Cell Technologies 100-0276) every other day and passaged every 3–4 days with Accutase (Stem Cell Technologies 07920) or ReLeSR (Stem Cell Technologies 100-0483). After passaging, iPSCs were plated into mTeSR Plus with 10 µM Y- 27632 (SelleckChem S1049).

### hNIL engineering

Patient-derived N98S-P2, E396K-P1, and E396K-P2 iPSC lines were engineered to contain a doxycycline inducible vector expressing human transcription factors NGN2, ISL1, and LHX3 (hNIL) in the CLYBL safe harbor locus as previously described ^4,26^. Briefly, after nucleofection of the hNIL vector and CLYBL-targeting TALEN plasmids, red fluorescent iPSC colonies were isolated and genotyped via junction PCR to verify hNIL integration in the CLYBL locus, and via copy number ddPCR. The ddPCR assay used a custom TRE3G or Neomycin copy number assay with RPP30 copy number assay (Bio-Rad dHsaCP1000485) as a 2-copy control. Clones with homozygous hNIL integration were selected and mCherry-neomycin resistance selection cassette was removed via nucleofection of a Cre recombinase plasmid (Addgene plasmid #11543). Post-transfection, non-fluorescent clones were further genotyped via junction PCR (Table S10) and copy number ddPCR using a custom TRE3G copy number assay and RPP30 ddPCR copy number assay (Table S9).

### RNP transfections

To prepare the ribonucleoprotein (RNP) for single-guide editing, 240 pmol of the gRNA (Synthego) was mixed with 120 pmol of Hi-Fi SpCas9 protein (HiFiCas9, Macrolab) and incubated for 30 minutes at room temperature. For dual-gRNA excision, 120 pmol of each gRNA (Synthego) was complexed separately with 60 pmol of HiFiCas9 protein and incubated for 30 minutes at room temperature. The RNP complexes were combined immediately prior to transfection. All gRNA sequences are listed in Table S2. After dissociation with Accutase, 3.0 x 10^5^ cells were resuspended in P3 buffer containing the RNP complex(es). Optionally, 50 pmol of ssODN was added to the cell suspension at this step. All ssODN sequences are listed in Table S3. iPSCs were nucleofected using the P3 Primary Cell 4D-Nucleofector X Kit S (Lonza, V4XP- 3032) with pulse code DS138. After nucleofection, cells were incubated for 5 minutes at room temperature and then plated into mTeSR Plus with 10 µM Y-27632. Genomic DNA was extracted from edited and unedited cells 3-4 days post-nucleofection using the DNeasy Blood and Tissue Kit.

### Isolation of clonal edited lines

The N98S-P2 fs and N98S-P2 cor iPSC lines were generated and maintained as previously described^4,45^, with minor changes to the maintenance conditions (i.e. mTeSR Plus was used in place of Stemfit). To generate excision and inversion iPSC lines, iPSCs were nucleofected with paired RNP complexes as above. After nucleofection, the cells were seeded in a 6-well plate with serial dilution to obtain a range of seeding densities across the plate. Cells were maintained for 7 days or until individual iPSC colonies were large enough for manual clone picking. Individual colonies were picked and transferred into a 48-well plate coated with Matrigel (Corning 356231). Once confluent, they were further expanded and gDNA was harvested using Quick Extract DNA Extract Solution (Lucigen, QE9050). Excision and inversion clones were genotyped via ddPCR and Sanger sequencing (Table S66).

### i^3^LMN differentiation

Patient-derived iPSC lines engineered with the hNIL transgene were differentiated into i^3^LMNs as described^4,26^. For gene expression assays, 1.25 x 10^5^ cells per cm^2^ were seeded onto poly-D-lysine (PDL, Sigma, P7405) and laminin-coated 6-well plates on day 3. For imaging assays, approximately 5-8 x 10^4^ cells per cm^2^ were seeded into similarly coated 96-well plates on day 3.

### In vitro cutting assay

RNP was complexed by addition of 1 μl of 10 uM Cas9 with 3 μl of 10 uM gRNA in 3 μl NebBuffer r3.1 (NEB B6003S) and 20 μl DNAse/RNase free water. This was incubated at room temperature for 10 minutes. Purified PCR amplicon (3 μl of 125 ng/ μl) was added to the reaction and vortexed lightly, followed by a 15-minute incubation at 37 °C. Proteinase K (1 μl) was added, and the reaction was incubated at room temperature for 10 minutes. The reaction was run via gel electrophoresis on a 2% agarose gel and stained with SYBR Safe for imaging.

### Measuring indels with amplicon sequencing

To measure single guide editing efficiency and specificity via next-generation sequencing (NGS), the targeted regions were amplified with primers containing Illumina adapter sequences (Table S11). PCR amplicons were isolated using PCR cleanup beads from the UC Berkeley DNA Sequencing Facility and eluted in DEPC-Treated water. Samples were submitted to Quintara Biosciences for Illumina-based NGS. FASTQ files were processed with Geneious Prime and the editing outcomes analyzed using the allele specific CRISPResso2 pipeline^33^. For Sanger sequencing, PCR amplicons were submitted to Quintara Biosciences and sequencing traces were analyzed using Synthego’s Inference of CRISPR Edit (ICE) Tool^46^.

### Measuring excisions and inversions

Excision and inversion frequency were measured via droplet digital PCR (ddPCR) according to the methods previously described using the BioRad QX200 System and QuantaSoft software^17^. Briefly, for excision assays forward and reverse primers flanked the two gRNA target sites. A FAM-labeled probe (IDT) was designed to bind within the PCR amplicon but outside the predicted excision. For inversion assays, the same FAM probe and its adjacent primer were used with a primer oriented between the two gRNA cut sites such that amplification could occur only in the presence of inversion. Each excision/inversion reaction included a HEX-labeled RPP30 copy number assay (Bio-Rad, dHsaCP2500350) as an internal reference for normalization.

### ddPCR specificity assay for excision and inversion

A multi-step PCR-ddPCR assay was designed to quantify the percentage of excision or inversion edits that occurred on each allele, as illustrated in Figure S15. Primers were designed to amplify the excised or inverted DNA including the SNP rs2979685, which is upstream of the *NEFL* gene. PCR amplicons were run on a 1% agarose gel and a band of the expected size was extracted using QIAquick Gel Extraction Kit (Qiagen, 28706). Purified amplicon DNA (0.01 pg) was used in a 25 μl ddPCR reaction containing a TaqMan SNP genotyping assay targeting rs2979685 (Thermofisher, Assay ID: C 11857476_30). Fractional abundance of the FAM vs. VIC signal determined the percentage of target allele versus non-target allele excised or inverted. All primer and probe combinations are listed in Table S7 with their sequences in Table S8.

### dPCR assay for combined excision frequency and specificity

A four-color assay using the QIAcuity One 5plex Device (Qiagen 911021) was used to measure excision frequency and specificity by single-step digital PCR as illustrated in Figure S16.

Genomic DNA (25–150 ng) was used in a 12 μl reaction containing a Taqman SNP genotyping assay for rs2979685 (FAM/VIC), a custom-designed primer and ROX-labeled probe mix to detect excision events, and Cy5-labeled RPP30 copy number control (Bio-Rad, dHsaCNS753942929). The reactions were loaded into an 8.5k partition 96-well plate and processed in a QIAcuity One instrument. The QIAcuity Software Suite was used for data analysis as follows and illustrated in Figure S16. Excision frequency was calculated as the ratio of ROX/Cy5 positive partitions. Specificity was calculated by the number of partitions positive for both excision (ROX) and the target allele (FAM or VIC) divided by the total number of double-positive partitions (ROX+FAM or ROX +VIC). Ambiguous partitions that were triple-positive (ROX+FAM+VIC) were excluded from the calculation but consistently represented less than 10% of all ROX-positive partitions.

### Off-target analysis

Sanger sequencing was used to determine if indels were produced at predicted off-target sites in the CRISPR-engineered cell lines as previously described. In brief, targeted amplification and sequencing was performed for all genomic sites with a > 0.25 cutting frequency determination (CFD score)^48^. This analysis was performed for unedited N98S-P2 and E396K-P2, N98S-P2-fs, N98S-P2-cor, N98S-P2-ex, N98S-P2-inv, E369K-P2-ex, and E369K-P2-inv. All sequences that deviated from reference were present in both the edited and unedited controls, consistent with pre-existing variants in the parental lines.

### Gene expression analysis by ddPCR

RNA was extracted from i^3^LMNs using the Quick-RNA Miniprep Kit (Zymo Research R1055), according to the manufacturer’s instructions. RNA concentrations were quantified using the NanoDrop spectrophotometer. Reverse transcription was performed using the iScript cDNA Synthesis Kit (Bio-Rad 1708891). To determine total mRNA expression, i^3^LMN cDNA (0.5 ng) was amplified in a 25 μl ddPCR reaction using commercially available FAM-labeled TaqMan gene expression assays for *NEFL* (ThermoFisher, Hs00196245_m1) or *NEFM* (ThermoFisher, Hs00193572_m1) in combination with HEX-labeled *GAPDH* gene expression assay (BioRad 10031255) as an internal control for normalization. To determine allele-specific *NEFL* mRNA expression, i^3^LMN cDNA (0.5 ng) was amplified in a 25 μl ddPCR reaction with TaqMan SNP genotyping assay targeted rs79736124 SNP in the 3’ UTR of *NEFL* (ThermoFisher, C_105316276_10). All ddPCR reactions were analyzed using the BioRad QX200 System and QuantaSoft software. For total mRNA expression of *NEFL* and *NEFM*, the ratio of FAM- positive to HEX-positive droplets was calculated to normalize to *GAPDH*. Fractional abundance of FAM-positive vs. VIC-positive droplets was used to quantify allele-specific *NEFL* expression.

### Immunofluorescent staining

Immunofluorescent staining was conducted as previously described^4^ with the following modifications: i^3^LMNs were cultured in clear-bottom imaging 96-well plates and fixed with an equivalent volume of 4% paraformaldehyde (PFA, Electron Microscopy Sciences 50-980-487) in PBS added directly to the cell culture media. To minimize lifting of the neuronal culture, the plate was held upright so that the media pools at the bottom of the well; using a multichannel pipette, media was slowly skimmed off the top of the pool and added slowly to the edge of the pool. Excessive drying of the wells was avoided. Cells were incubated in PFA at room temperature for 20 min, then washed with PBS with 0.1% Triton-X (PBS-T, Sigma X100) once quickly, followed by two 15-minute washes. The cells were blocked and permeabilized for 1 hour at room temperature with 5% Normal Goat Serum (NGS, ThermoFisher Scientific PCN5000) in PBS-T. The cells were incubated in primary antibodies at appropriate dilutions (Table S6) in 3% NGS in PBS-T at 4 °C overnight. The following day, the cells were washed with PBS at room temperature once quickly, followed by three 10 minute washes. The cells were incubated in fluorescent conjugated secondary antibodies diluted at 1:500 in 3% NGS in PBS-T for 1 hour at room temperature. The cells were washed with PBS once quickly, followed by three 10-minute washes. The first 10-minute wash contained DAPI dye (Invitrogen, D1306).

### Microscopy and imaging analysis

Images were taken on a Keyence BZ-9000 Fluorescence Microscope with 40x objective or the automated Cell Insight CX7 LZR Pro HCS Platform at 20x (CX7, ThermoFisher Scientific). The CX7 microscope has a field size of 444.54 by 444.54 microns and the camera acquisition mode is 1104x1104 (2x2 binning). To maximize the consistency and reproducibility of this analysis, all neurons for each experiment were fixed and stained simultaneously using a single aliquot and dilution of each antibody prior to imaging. Five (Keyence) or fifteen (CX7) images were acquired per well. All images for each experiment were acquired in a single session with constant illumination and exposure parameters.

The cell bodies and neurites of i^3^LMNs were segmented using CellProfiler pipelines as illustrated in Supplemental Figures S7 and S8. To measure fluorescence intensity in the cell body, first the CellProfiler pipeline defined the nuclei of neurons with HB9 (original) or with DAPI (modified). The diameter of each nucleus was expanded by 10 pixels to encompass the entirety of the cell body. After thresholding, this inferred cell body mask was overlaid onto the NF-L channel. Objects were filtered by compactness and shape to eliminate objects that are not cell bodies and mean intensity was measured. To measure fluorescence intensity in the neurites, the pipeline similarly identified DAPI-positive nuclei and expanded the objects by 15 pixels to encompass the cell body and axon hillock. NF-L and beta-tubulin channels were thresholded and an inverted mask of the inferred cell bodies was overlaid on the thresholded beta-tubulin channel to produce a neurite mask. The neurite mask was overlaid on the thresholded NF-L channel and total fluorescence intensity was measured and divided by the total area of the neurite mask to normalize for neurite density.

### Multi-electrode array

N98S-P2 and its associated frameshift, corrected, excision, inversion lines, along with E396K and its associated excision and inversion lines were differentiated into i^3^LMNs and seeded in 24- well CytoView MEA plates (Axion Biosystems, M384-tMEA-24W) on day 3 as previously described^4^. BrainPhys (StemCell, 05791) was used as basal media starting on day 7. Spontaneous neural activity was measured for 15 minutes on days 7, 14, 21 and 28 using the Axion Biosystem Maestro Edge MEA systems and analyzed as previously described^4^.

### Statistics

All statistical analyses were performed using GraphPad Prism (v10.4.1). Pairwise comparisons were done by unpaired two-tailed T-test. Multiple comparisons with a single variable were performed by one-way ANOVA followed by the recommended post-test as follows: Dunnett’s test for comparisons of multiple treatments to a control, Tukey’s test for comparisons of each treatment to every other treatment, and Šídák’s test for comparisons of pre-selected pairs of treatments. Comparisons of multiple variables were performed by two-way ANOVA followed by Šídák’s test for selected variables. Details of the parameters and outputs for the statistical tests used for each individual experiment are included in the supplemental materials.

### Computational methods

The frequency of heterozygotes in the 1000 Genomes Phase 3 dataset was used to select guide-pair candidates^28^. Common variants (allele frequencies between 0.1 and 0.9) were considered in the 50 kb regions on either side of *NEFL*. This resulted in a total of 151 SNPs: 66 upstream and 85 downstream of the *NEFL* gene body. Then for each flanking pair of SNPs the number of individuals out of the total of 2,548 that were heterozygous for both SNPs was identified. To explore the linkage disequilibrium patterns in the region, these 151 SNs, plus 3 common (0.1 < allele frequency < 0.9) SNPs from within the *NEFL* gene, were computed across all 1000 Genomes populations and visualized using the LDMatrix tool on the NIH LDLink website (https://ldlink.nih.gov/?tab=ldmatrix)^44^.

To identify a small collection of SNP pairs that enable editing a large number of diverse individuals, a greedy algorithm was applied to select SNP pairs. At each iteration, the pair of SNPs was chosen that was heterozygous in the largest number of individuals not covered by a previously selected pair. Those individuals were then removed from consideration, and the process was repeated until an acceptable proportion of individuals was covered or a maximum number of SNP pairs was reached.

1000 Genomes variants (in VCF format) are available at (ftp.1000genomes.ebi.ac.uk/vol1/ftp/data_collections/1000_genomes_project/release/20190312_ biallelic_SNP_and_INDEL/)^28^.

## Supporting information

Statistics supplemental figures

Statistics main figures

## Author Contributions

LMJ and BRC conceived the project. PHD, BMJS, CBEM, HLW, and LMJ designed the experiments. PHD, CMF, CBEM, BMJS, and HLW completed cell line generation and characterization. PHD, CMF, CBEM, and BMJS performed gene editing experiments and assays. PHD and BMJS performed gene expression analysis. PHD, CBEM, and BJMS performed neuron differentiations and neuron imaging. PHD and CBEM performed multielectrode array analysis. CBEM, PHD, and DS designed and executed the CellProfiler analysis pipelines. AA, ENG, GDR, and JAC performed computational analyses. PHD, BMJS, and LMJ prepared the figures and wrote the manuscript with assistance from all authors.

## Acknowledgements

We thank Michael Shy for providing primary dermal fibroblasts from patients with E396K mutations. We thank Wendy Runyon, Hana Ghanim, and Angela Liu for technical assistance with iPSC reprogramming and integration of hNIL transgenes. We thank Anke Meyer-Franke and the Gladstone Assay Development and Drug Discovery Core for advice and technical assistance with automated microscopy. We thank the Gladstone Stem Cell Core, Bioinformatics Core, and Histology and Light Microscopy Core for their resources and services. We thank Bria Macklin, Zachary Nevin, Gokul Ramadoss, and Françoise Chanut for helpful discussions and review of the manuscript.

## SUPPLEMENTARY DATA

**Figure S1:**
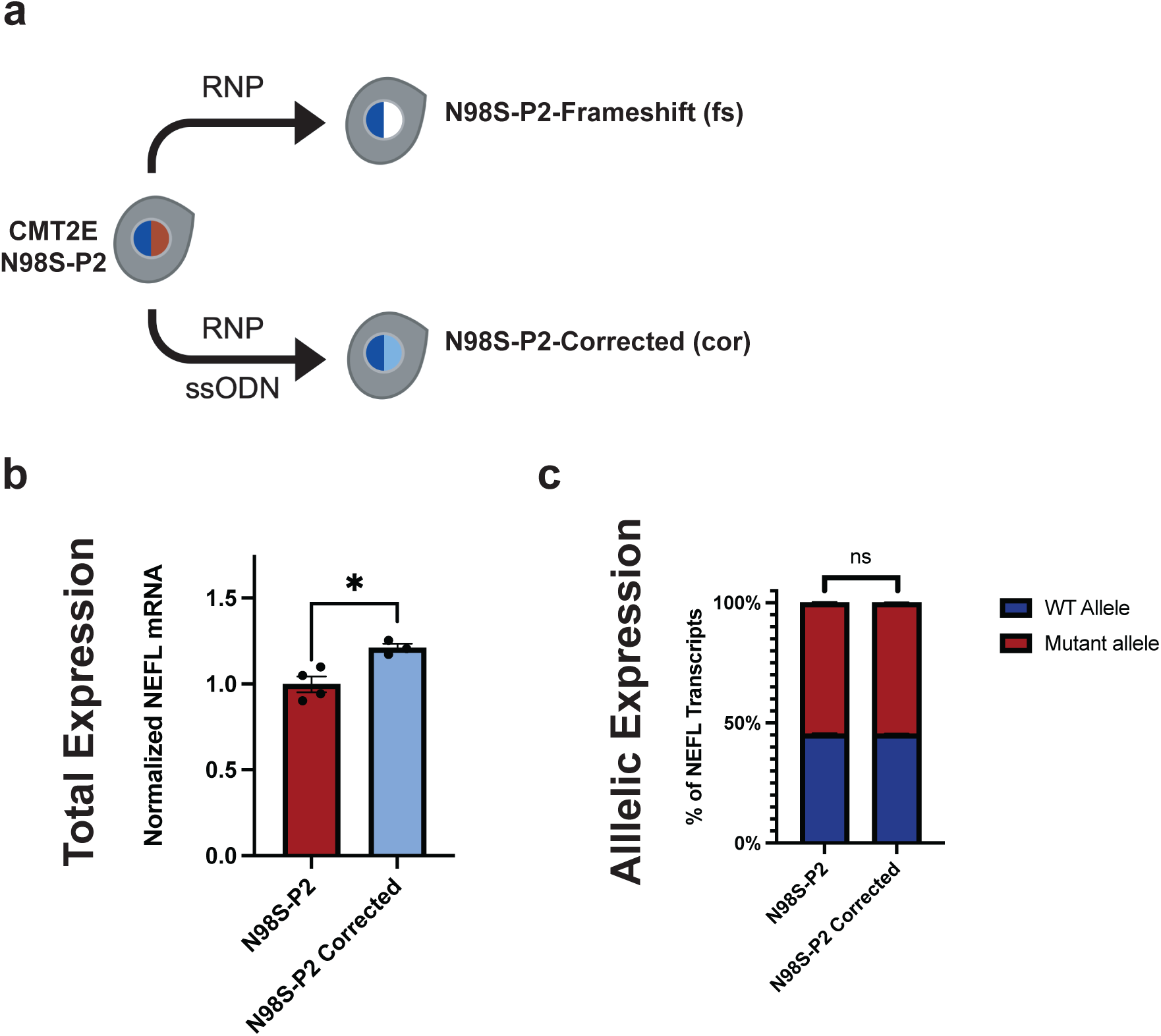
Derivation of clonal edited iPSC lines from the CMT2E N98S-P2 patient line. **(a)** Transfection of N98S-P2 iPSC with N98S-specific HiFiCas9 RNP produced N98S-P2-fs (contains +1 indel at the mutation locus). Transfection of N98S-P2 iPSC with N98S-specific HiFiCas9 RNP plus single-strand oligonucleotide donor produced N98S-P2-cor (precise correction of N98S mutation with linked silent mutation to facilitate genotyping). Colored nuclei indicate *NEFL* genotype: dark blue = wild type, red = N98S mutant, white = frameshift (knockout), light blue = N98S-corrected (+ silent mutation). N98S-P2 and N98S-P2-cor were differentiated into i^3^LMNs and RNA was extracted on day 7. **(b)** Total *NEFL* expression by quantitative RT-ddPCR, normalized to GAPDH. **(c)** Relative allelic expression by allele discrimination RT-ddPCR. Bar graphs represent mean +/- S.E.M. of biological replicates. * p < 0.05, ns p > 0.05.

**Figure S2:**
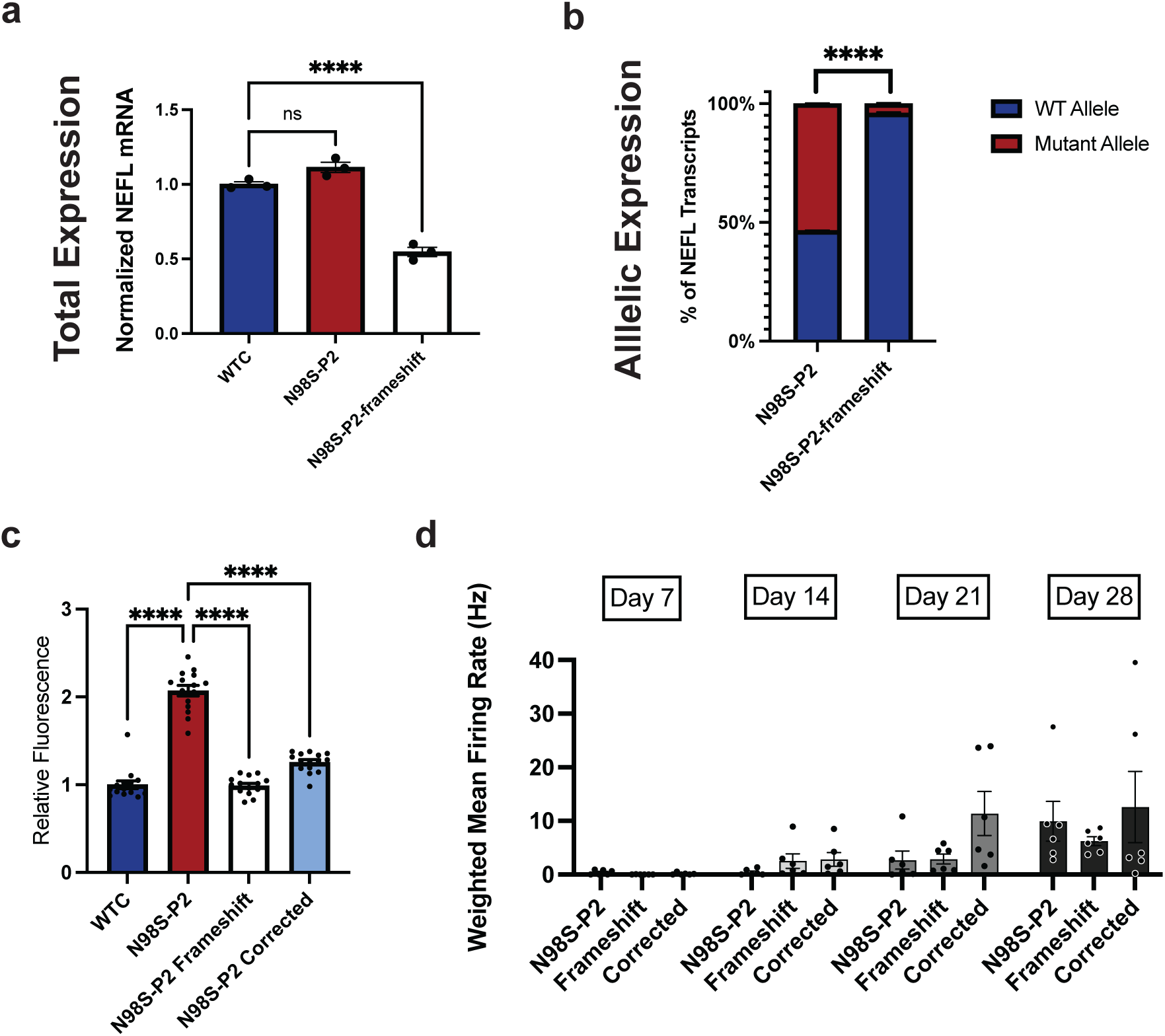
Frameshift editing in N98S-P2 i^3^LMNs inactivates the mutant allele and rescues disease phenotype. N98S-P2 and N98S-P2-fs were differentiated into i^3^LMNs and subsequently used for RNA extraction or immunofluorescent staining on day 7. **(a)** Total *NEFL* expression by quantitative RT-ddPCR, normalized to *GAPDH*. **(b)** Relative allelic expression by allele discrimination RT-ddPCR. **(c)** Mean NF-L fluorescence intensity in the cell bodies of iPSC-derived motor neurons at day 7. **(d)** Multi-electrode array was conducted to measure spontaneous action potentials in N98S-P2, and in corrected and frameshift lines. Spontaneous electrical activity was measured at days 7, 14, 21, and 28. Each data point represents the mean weighted firing rate from one biological replicate. Two-way ANOVA demonstrated a statistically significant effect of the differentiation day (p < 0.001) but no significant difference between cell lines (p > 0.05). Bar graphs represent mean +/- S.E.M. of biological replicates. **** = p < 0.0001, ns p > 0.05.

**Figure S3:**
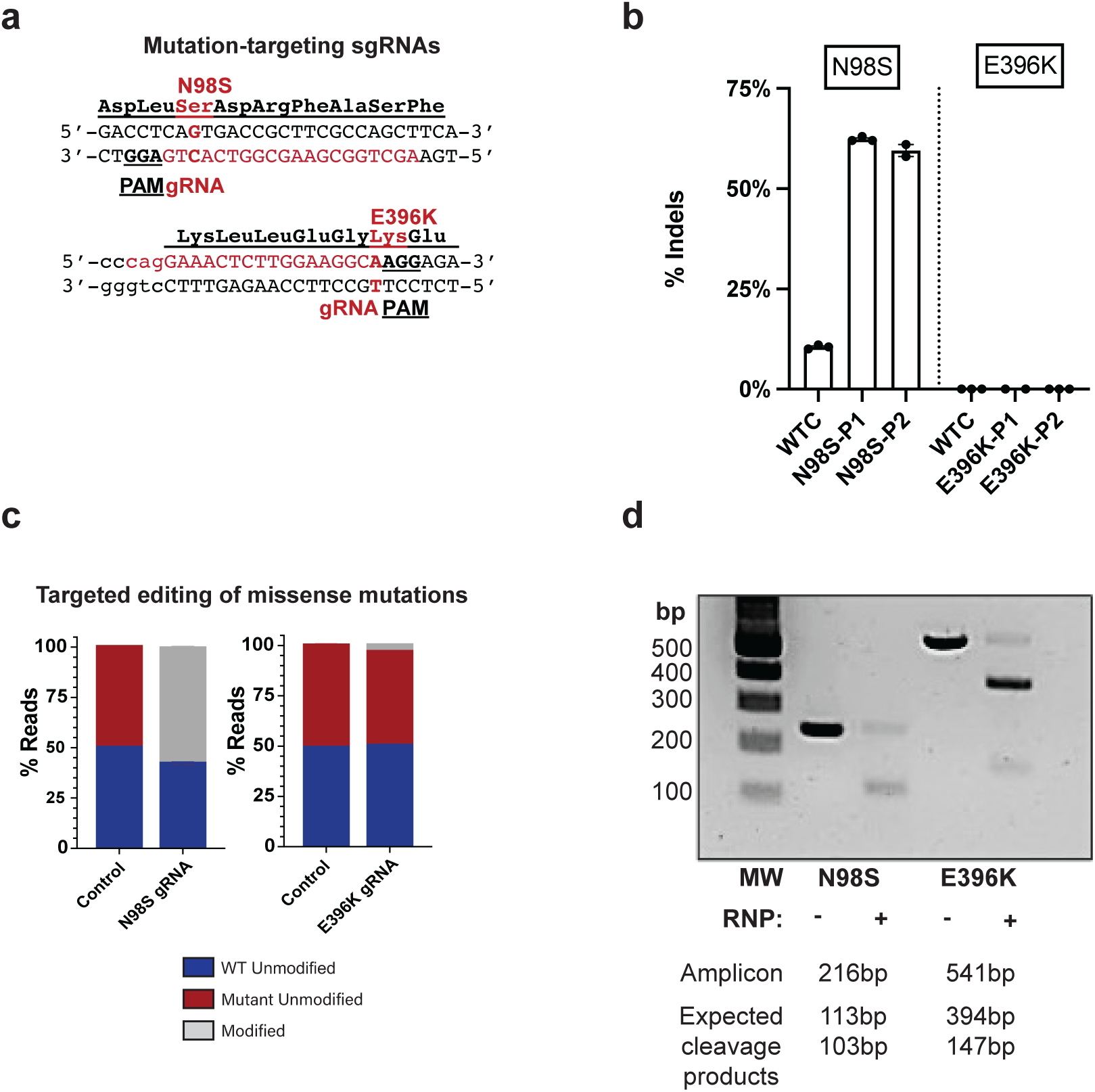
Comparison of the editing and nuclease activity of gRNAs targeting the N98S and E396K mutations. **(a)** Sequences of N98S and E396K mutation-targeting gRNAs (red) with associated PAM sequences (underlined). **(b)** Indel editing efficiency of N98S and E396K mutation-specific gRNAs in multiple patient backgrounds by Sanger sequencing and ICE analysis. Bar graphs represent mean +/- S.E.M of replicate transfections. **(c)** NGS amplicon sequencing of N98S and E396K targeted editing in N98S-P2 and E396K-P2 iPSCs. Modified reads contain indels at the target site, WT and mutant unmodified indicate reads without indels on the WT and mutant alleles, respectively. **(d)** PCR amplicons spanning the N98S or E396K mutations were generated from N98S-P2 and E396K-P2 gDNA, respectively, followed by incubation with corresponding N98S or E396K mutation-specific gRNA and HiFiCas9 RNP. Gel electrophoresis identifies the expected cleavage products. MW = molecular weight DNA ladder.

**Figure S4:**
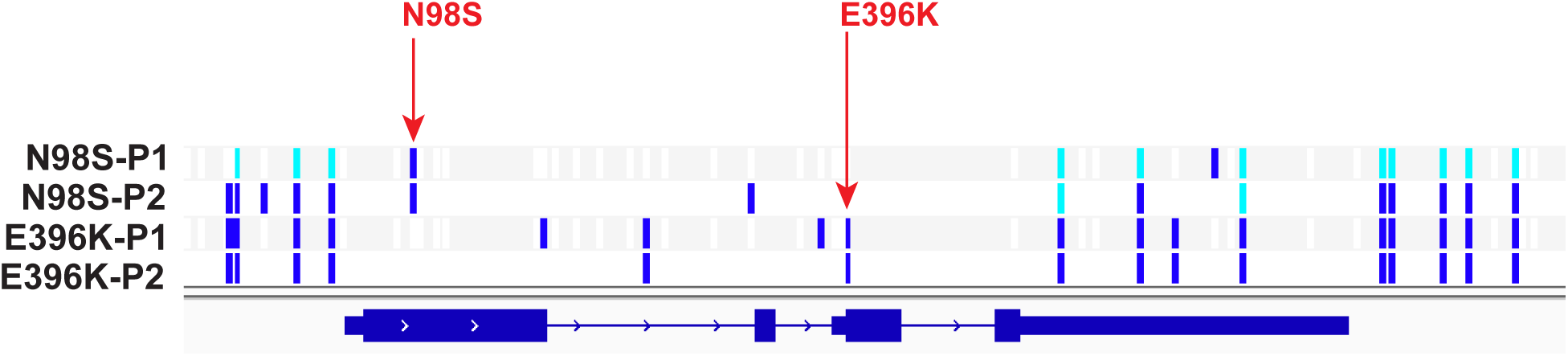
Panel of patient cell lines with variants flanking the *NEFL* coding region. Whole genome sequencing was performed from patient-derived iPSC. Dark blue lines indicate heterozygous variants, light blue lines indicate homozygous variants. Causative missense mutations are annotated with red arrows. Images generated from vcf files using Integrative Genomics Viewer^47^.

**Figure S5:**
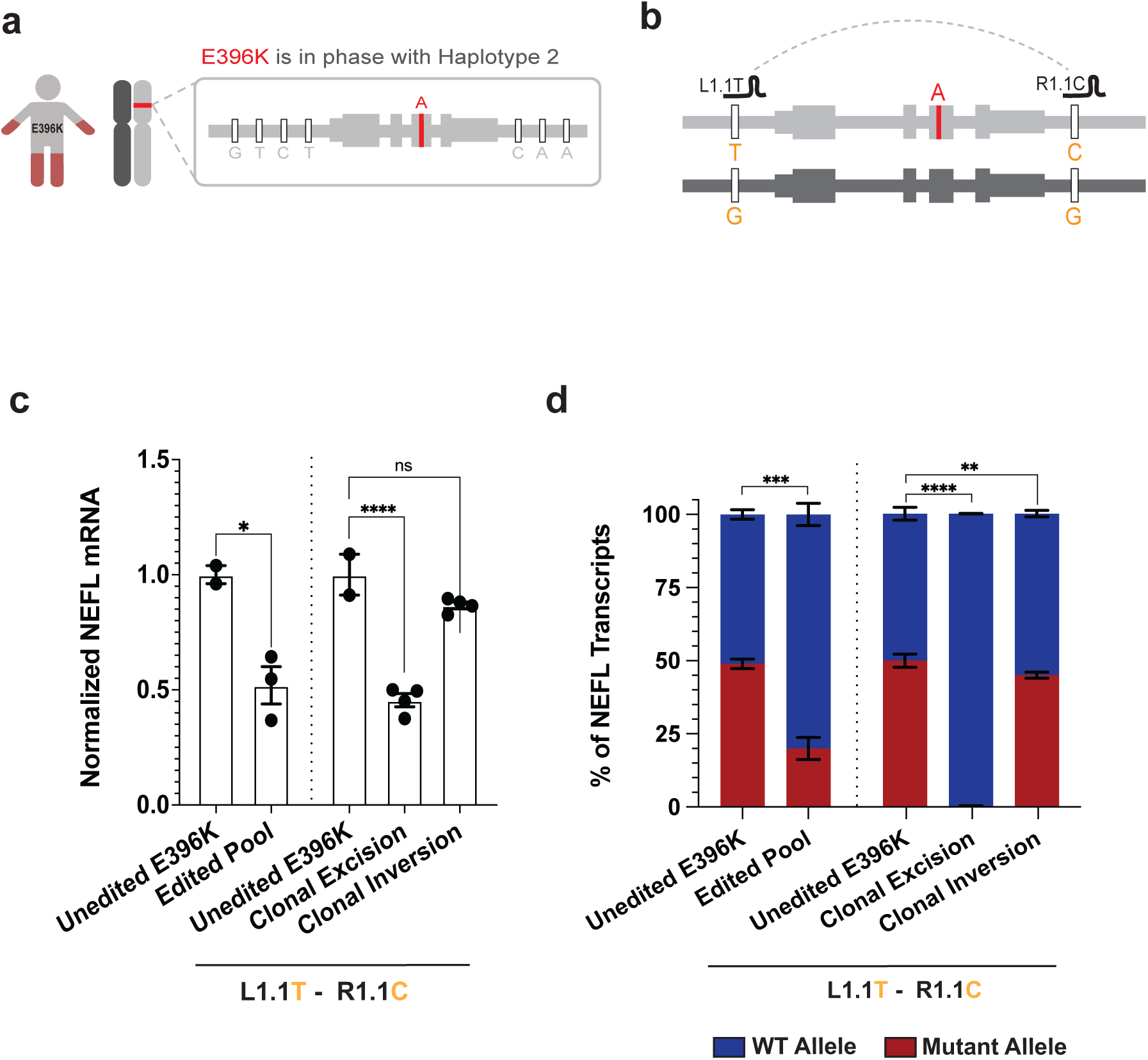
Analysis of *NEFL* gene expression after haplotype editing in E396K-P2 i^3^LMNs. **(a)** Schematic of phasing of E39K mutation with Haplotype 2 **(b)** Schematic of SNP-targeting guide pair for E396K haplotype editing. Edited and unedited iPSCs from E396K-P2 were differentiated into i^3^LMNs, and RNA was extracted on Day 7. **(c)** Total *NEFL* expression was measured by quantitative RT-ddPCR relative to *GAPDH* and normalized to the unedited control in. **(d)** Relative expression of wildtype and mutant alleles was measured by allele discrimination ddPCR. Bar graphs represent mean +/- S.E.M of biological replicates. * p < 0.05, ** p < 0.01, *** p < 0.001, **** p < 0.0001, ns p > 0.05.

**Figure S6:**
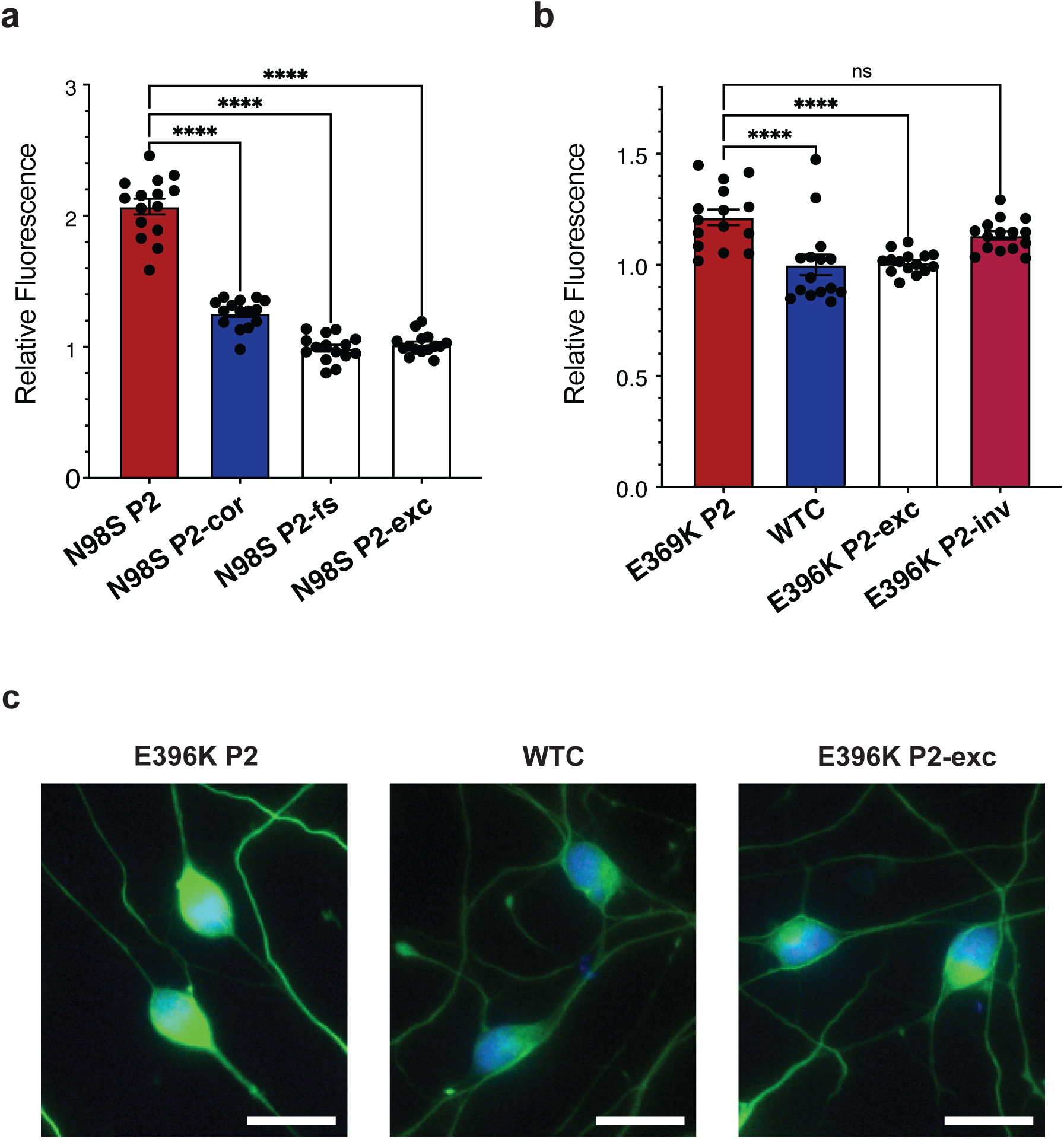
NF-L accumulation in cell bodies of control and edited i^3^LMNs. Quantification of mean NF-L fluorescence intensity in inferred cell bodies in **(a)** N98S-P2 and **(b)** E396K-P2 i^3^LMNs at day 7. Each data point is the mean NF-L intensity calculated across five images per replicate. Bar graphs represent mean +/- S.E.M of biological replicates. **** p < 0.0001, ns p > 0.05. **(c)** Representative images of E396K-P2 i^3^LMNs stained with anti-NF-L (green) and anti- HB9 (blue). Scale bars = 20 µM.

**Figure S7:**
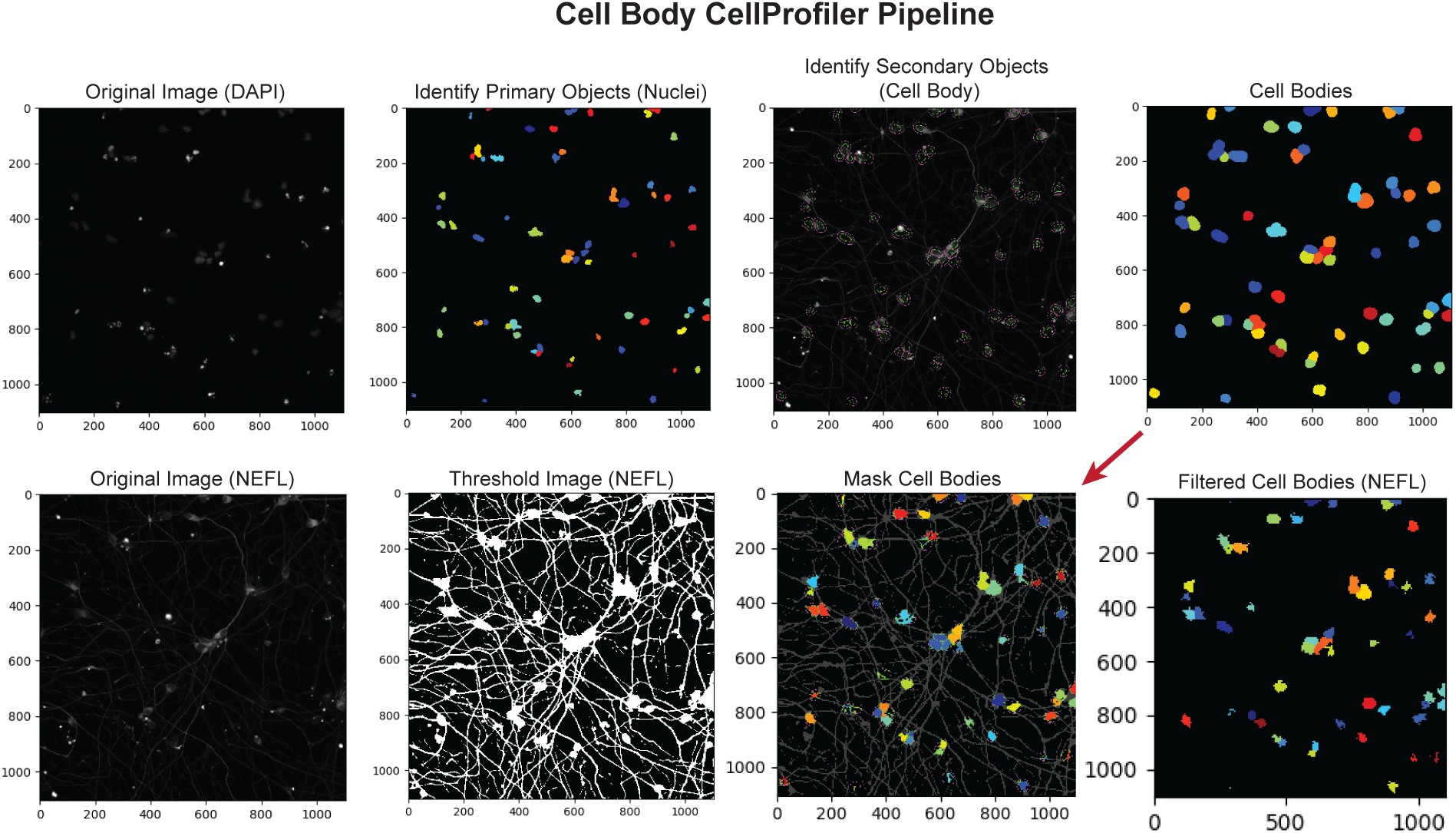
CellProfiler pipeline for measuring NF-L intensity in i^3^LMN cell bodies. Nuclei (primary objects) were identified from the DAPI-stained image channel. Cell bodies (secondary objects) were inferred by extending the area of DAPI+ signal by 10 pixels. The NF-L-stained image channel was thresholded and masked with these inferred cell bodies (red arrow). The cell bodies were then filtered by shape and compactness to remove any incorrectly identified objects. The mean intensity in the cell body-masked NF-L-stained image channel was measured to quantify NF-L within cell bodies.

**Figure S8:**
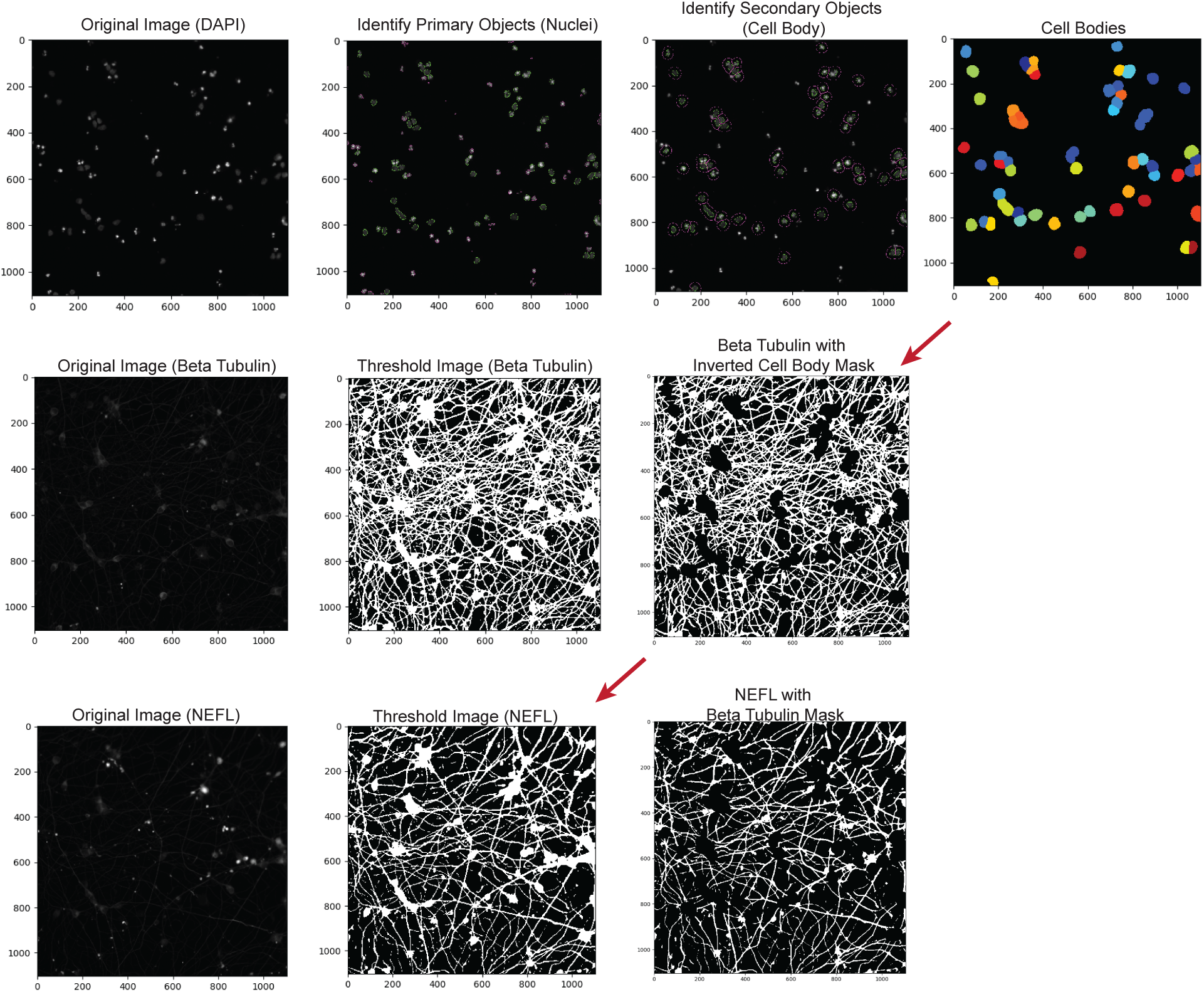
**CellProfiler pipeline for measuring NF-L intensity in i^3^LMN neurites.** Nuclei (primary objects) were identified from the DAPI-stained image channel. Cell bodies (secondary objects) were inferred by extending the area of the DAPI+ signal by 15 pixels. The beta3-tubulin-stained image channel was thresholded, and an inverted cell body mask was applied to exclude signal within the cell bodies and define the region encompassed by neurites. The NF-L-stained image channel was then thresholded and masked with the beta3-tubulin neurite image. Red arrows indicate the application of masks to images in other channels. Total intensity in the neurite-masked NF-L-stained image channel was calculated and divided by the beta3-tubulin-positive neurite area to normalize for variation in neurite density.

**Figure S9:**
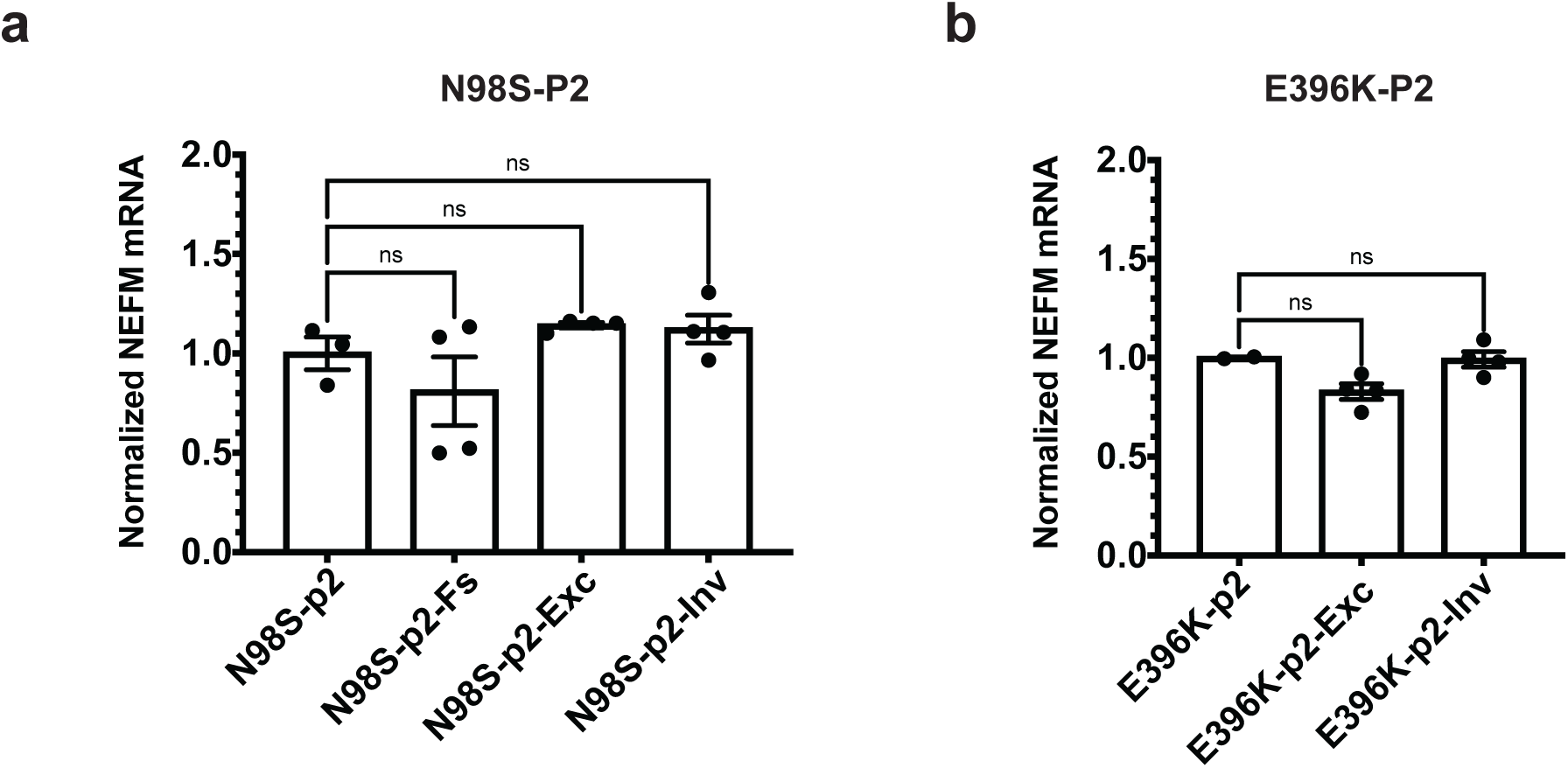
**Total *NEFM* expression in edited i^3^LMNs**. Edited iPSCs from **(a)** N98S-P2 and **(b)** E396K-P2 were differentiated into i^3^LMNs, and RNA was extracted on day 7. Total *NEFM* expression was measured by quantitative RT-ddPCR relative to *GAPDH* and normalized to unedited controls. Bar graphs represent mean +/- S.E.M. of biological replicates. ns p ≥ 0.05.

**Figure S10:**
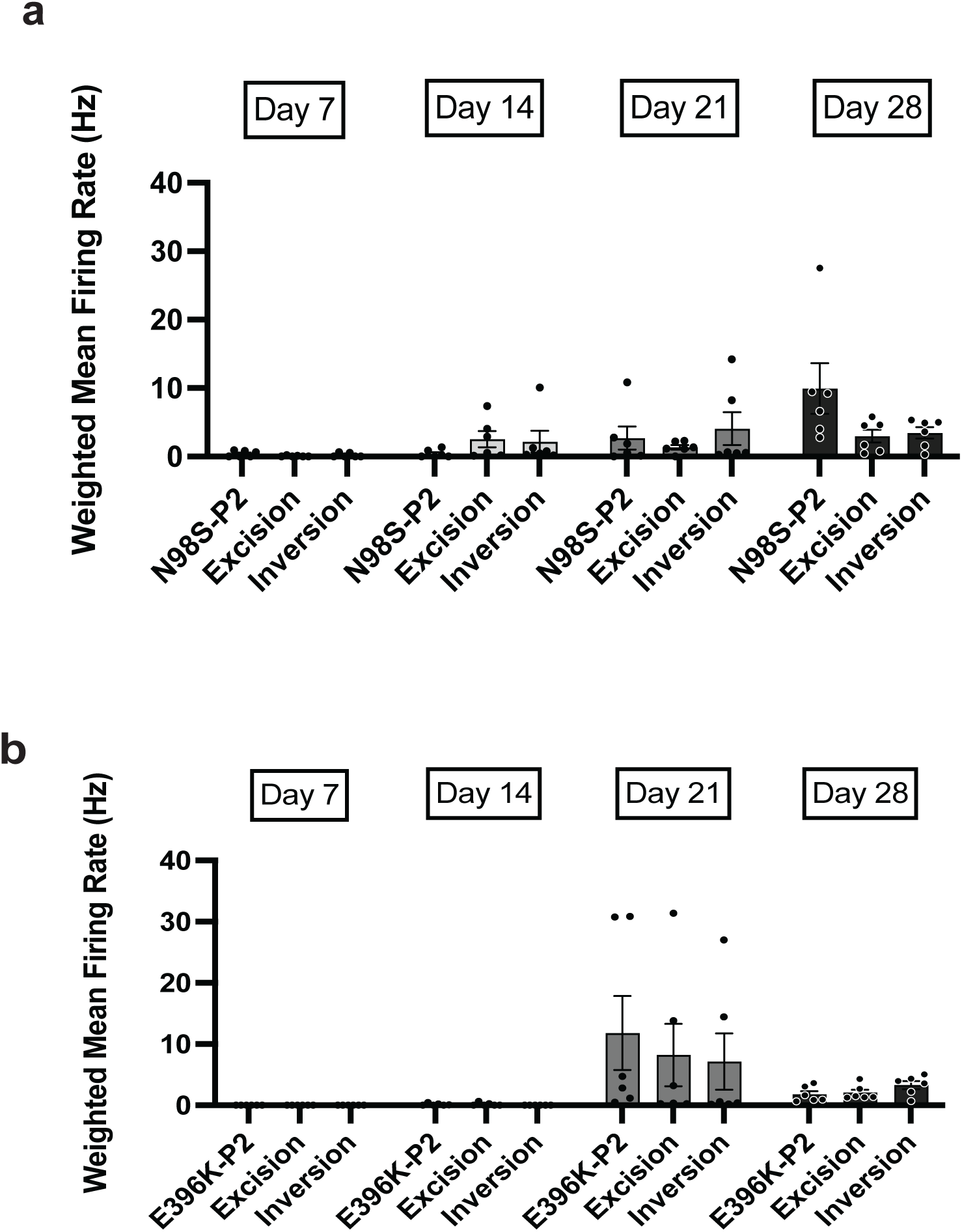
Total *NEFL* gene excision and inversion do not impact i^3^LMN electrophysiology. Multi-electrode array was conducted to measure spontaneous electrical activity at days 7, 14, 21, and 28 for **(a)** N98S-P2 and (b) E396K-P2 series of edited i^3^LMN. Each data point represents the mean weighted firing rate from one biological replicate. Two-way ANOVA demonstrated a statistically significant effect of the differentiation day (p < 0.001) but no significant difference between cell lines (p > 0.05). Bar graphs represent mean +/- S.E.M of biological replicates.

**Figure S11:**
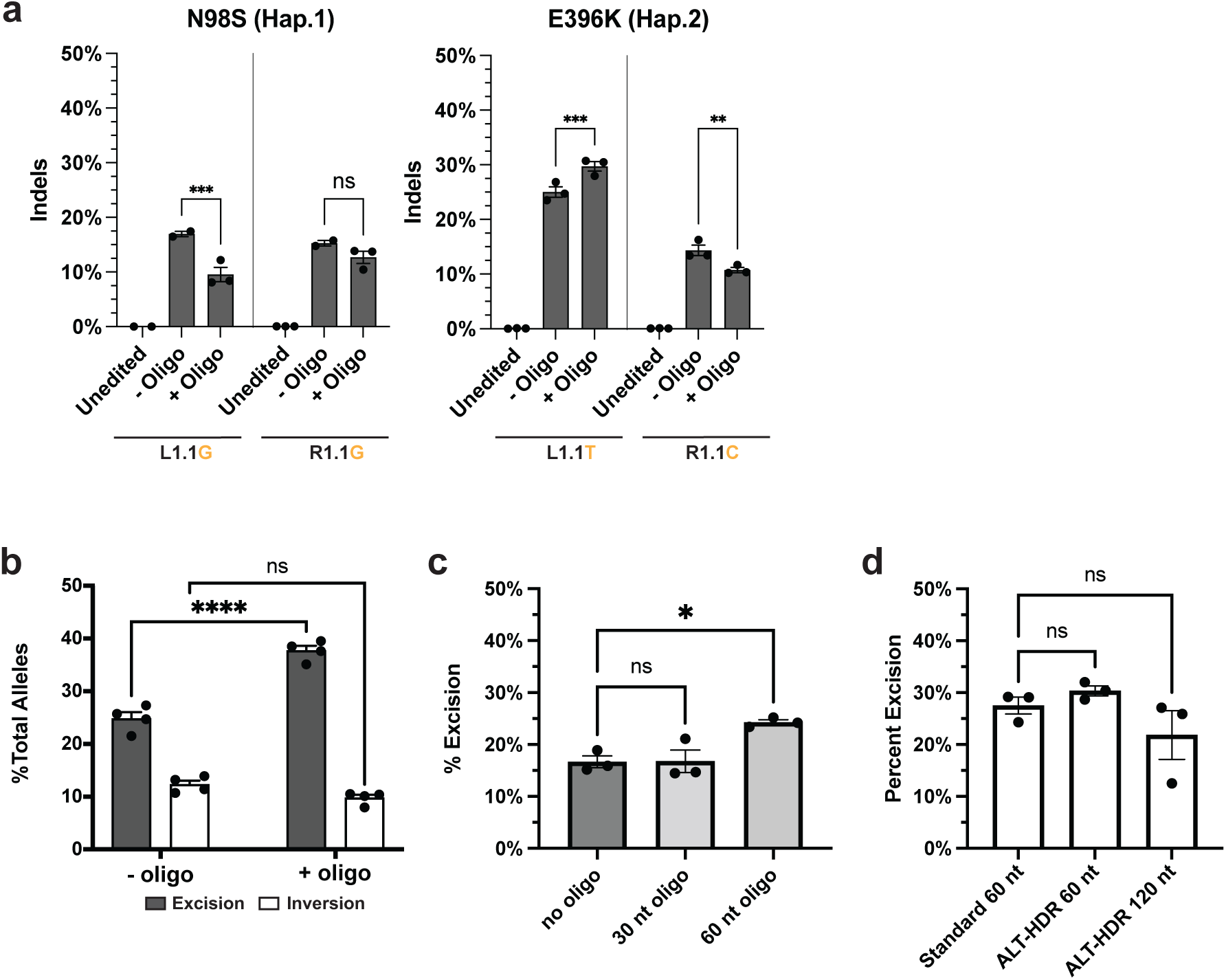
Effect of bridging oligos on editing outcomes. **(a)** N98S and E396K iPSCs were transfected with L1.1 and R1.1 RNP with and without the addition of a 60 nt single-strand oligonucleotide donor (ssODN). PCR amplicons spanning L1.1 and R1.1 target sites were analyzed by NGS amplicon sequencing to quantify indels on alleles that were not edited by excision or inversion. **(b)** N98S-P2 iPSCs were transfected with L1.1 + BA1 RNP with and without an excision bridging ssODN. Excision and inversion frequency were measured by ddPCR. **(c-d)** E396K-P2 iPSCs were transfected with L1.1 + R1.1 RNP and ssODN of different lengths and chemistries. Excision was measured by ddPCR. Comparison of 30 and 60 nt unmodified ssODN shown in (c). Comparison of unmodified (60 nt) and ALT-HDR modified (60 and 120 nt) ssODN shown in (d). For all experiments, gDNA was collected four days after nucleofection. Bar graphs represent mean +/- S.E.M. of replicate transfections. * p < 0.05, ** p < 0.01, *** p < 0.001, **** p < 0.0001, ns p > 0.05.

**Figure S12:**
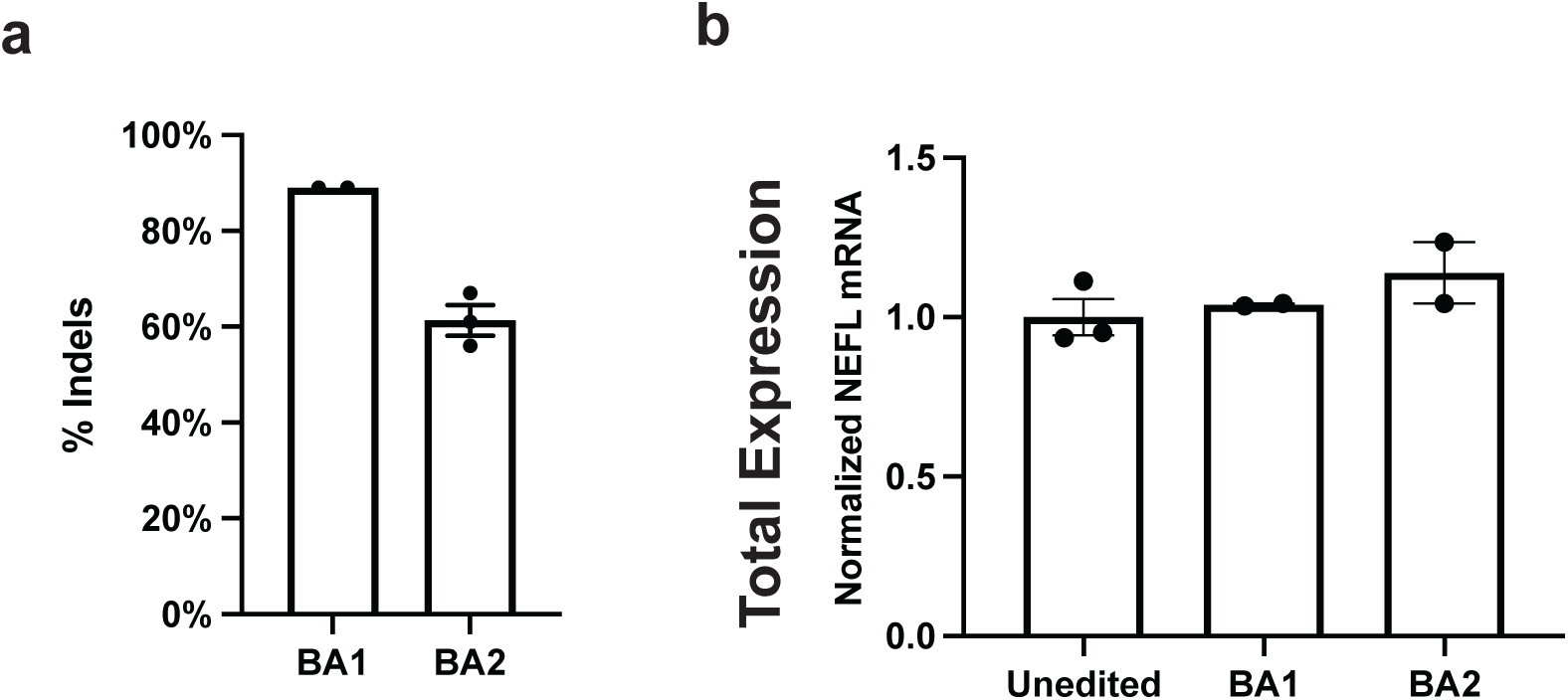
Evaluation of indels produced by biallelic gRNAs targeting *NEFL* intron 1. **(a)** Quantification of indels induced by BA1 and BA2 gRNAs as measured by Sanger sequencing and ICE analysis. **(b)** Total *NEFL* expression measured by quantitative RT-ddPCR, normalized to *GAPDH*. Bar graphs represent the mean of replicate transfections (a) or replicate differentiations (b) +/- S.E.M.

**Figure S13:**
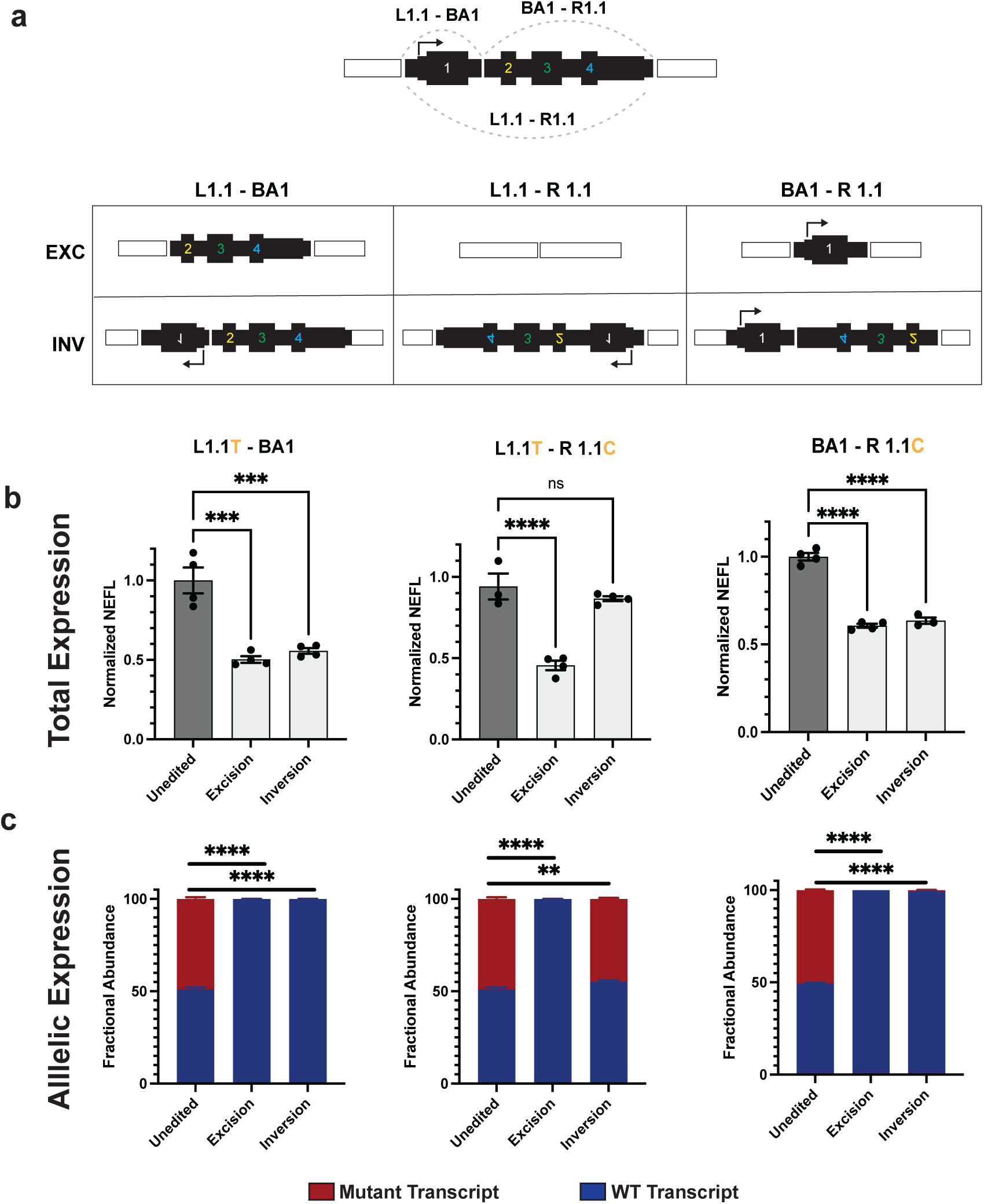
Clonal analysis of editing outcomes reveals variable contributions of excision and inversion to gene inactivation. **(a)** Schematic of predicted editing outcomes with various combinations of L1.1, R1.1 and intronic biallelic (BA1) gRNAs. E396K-P2 was transfected with each RNP pair and clonal iPSC lines were isolated with each predicted outcome and differentiated into i^3^LMNs. RNA was isolated on day 7 and measured by quantitative RT-ddPCR. **(b)** Total *NEFL* expression relative to *GAPDH* and normalized to unedited control. **(c)** Relative allelic expression via allele discrimination ddPCR using a heterozygous SNP in the 3’ UTR (rs2976439). Bar graphs represent the mean +/- S.E.M. of biological replicates. ** p < 0.01, *** p < 0.001, **** p < 0.0001, ns p > 0.05.

**Figure S14:**
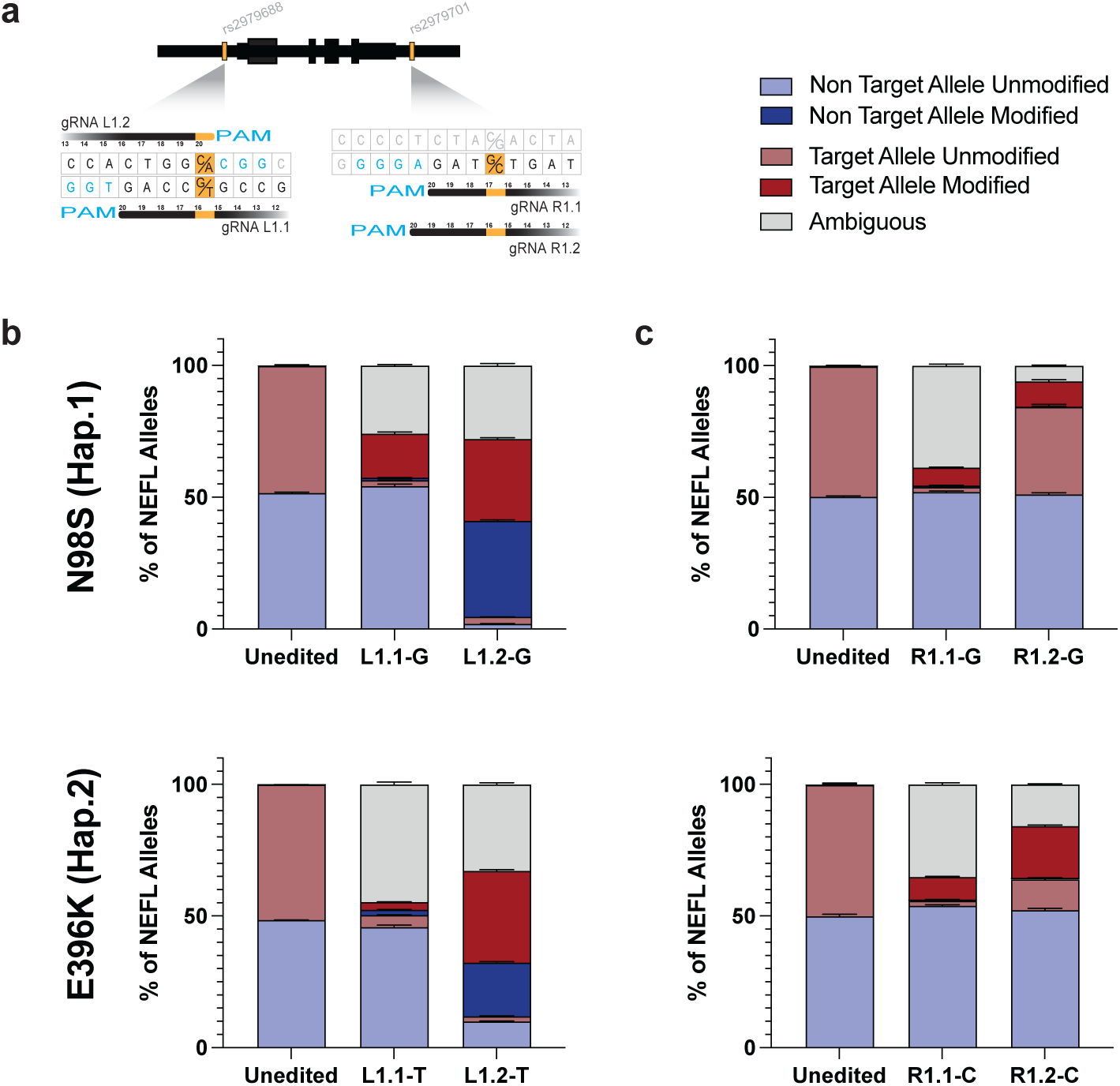
Analysis of allele-specific editing with single gRNA. **(a)** Schematic of sequences targeted by SNP-specific gRNAs with variants in orange and the associated PAM in blue. **(b-c)** N98S-P2 and E396K-P2 iPSC were transfected with the indicated allele-specific gRNA- HiFiCas9 RNPs followed by quantification of editing outcomes by NGS amplicon sequencing and CRISPResso2 analysis. Ambiguous editing events cannot be assigned to either allele due to the deletion of the variant nucleotide. Bar graphs represent the mean +/- S.E.M. of triplicate transfections.

**Figure S15:**
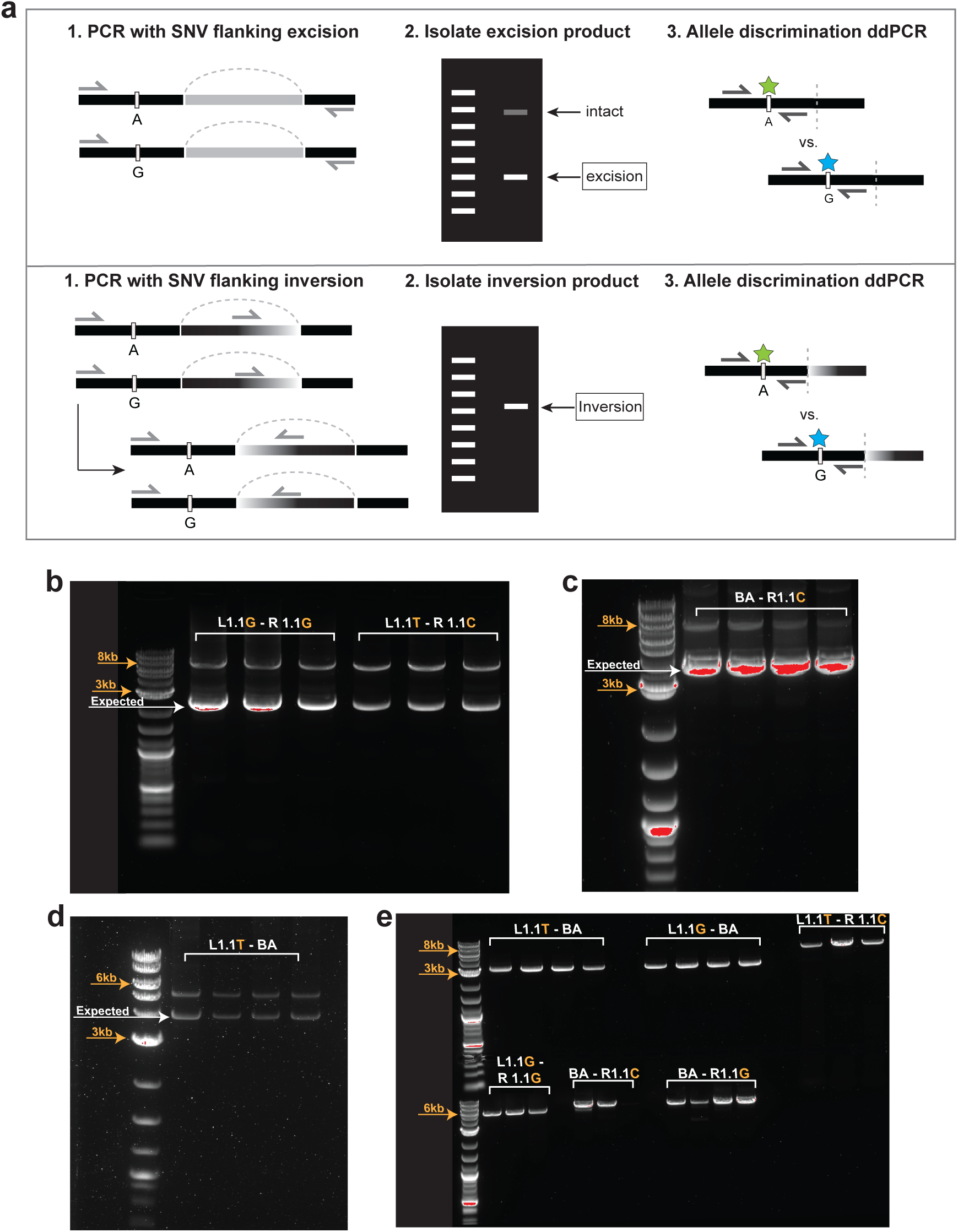
Multi-step PCR/ddPCR assay to measure allele-specificity of excision and inversion. **(a)** Assay workflow 1: Schematic of PCR designed to amplify excised or inverted alleles with flanking heterozygous variant; 2: Gel purification to isolate excision or inversion products 3: Schematic of allele discrimination ddPCR used to quantify specificity. **(b-d)** Examples of gel electrophoresis of PCR products used for excision specificity. **(e)** Examples of gel electrophoresis of PCR products used for inversion specificity.

**Figure S16:**
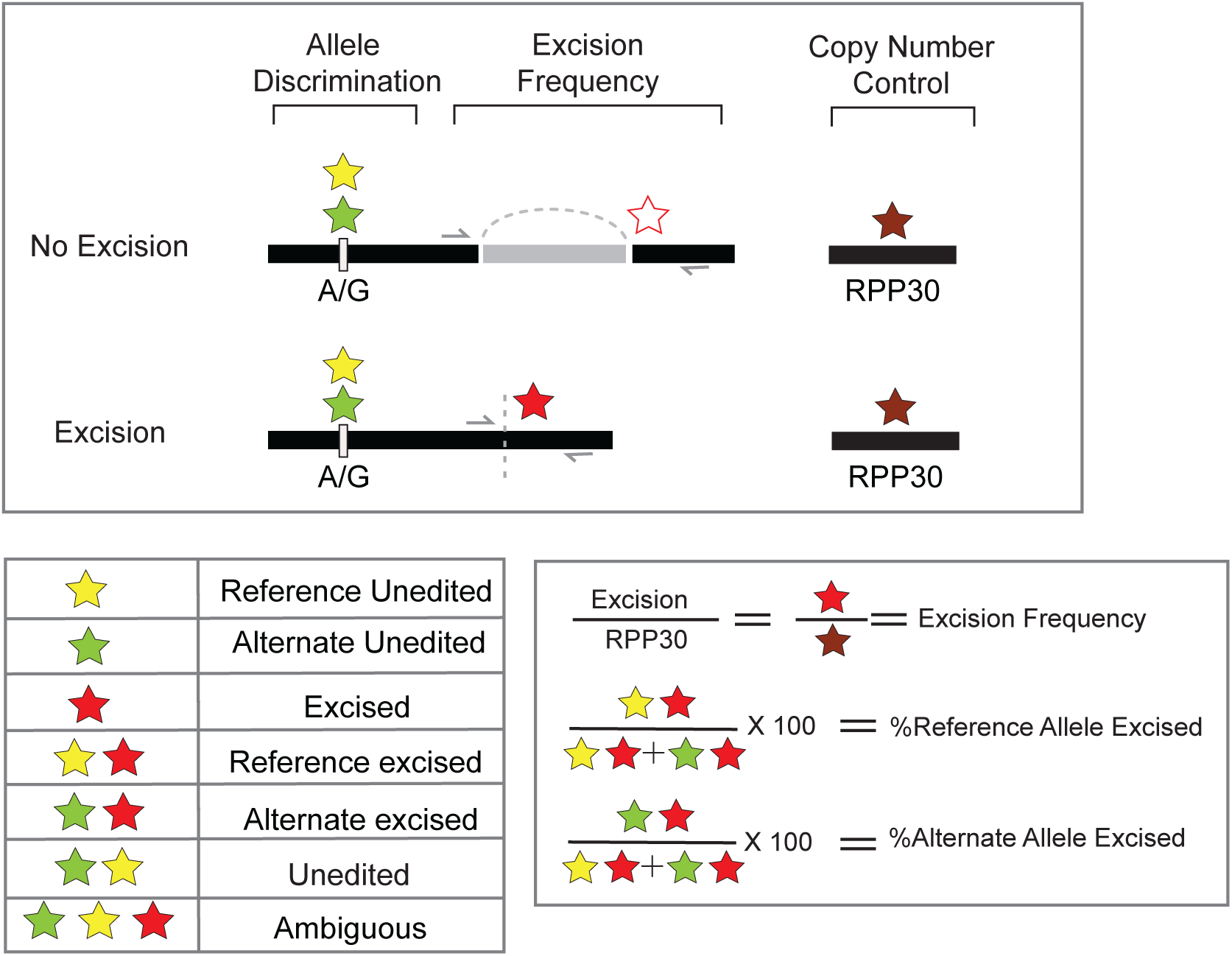
Single-step assay to measure excision frequency and specificity via multiplexed four-color digital PCR. Top: Schematic of multiplexed assay design with four probes (Green = FAM, Yellow = Hex, Red = ROX, Maroon = Cy5). **Bottom-left:** Partition occupancy observed for excision specificity measurements. **Bottom-right**: Calculation used to quantify excision specificity and frequency.

**Figure S17:**
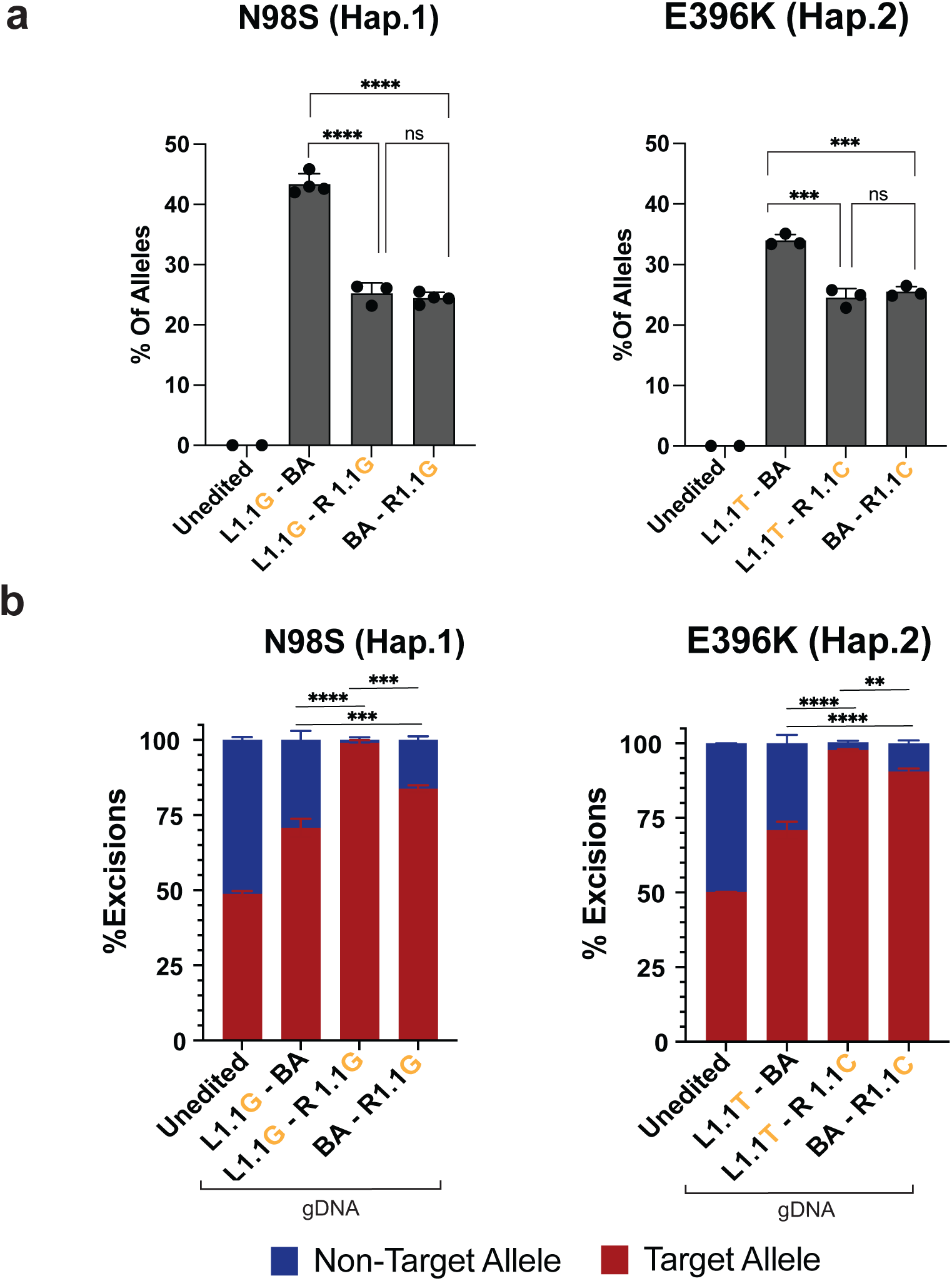
Quantification of excision frequency and specificity via multiplexed four-color digital PCR. **(a)** Excision frequency as measured by the ratio of excision (ROX+) signal over RRP30 (Cy5+) signal. **(b)** Excision specificity measured via dPCR allele-discrimination assay for a heterozygous SNP (rs2979685, ref = HEX, alt = FAM) located 5’ of *NEFL.* Unedited control represents the fractional abundance of HEX vs. FAM in ROX-negative partitions to demonstrate equal abundance of the two alleles at baseline, while edited samples represent the fractional abundance of HEX vs. FAM in ROX-positive partitions that contain an excision event. Bar graphs represent mean +/- S.E.M. of replicate transfections. ** p < 0.01, *** p < 0.001, **** p < 0.0001, ns p > 0.05.

**Figure S18:**
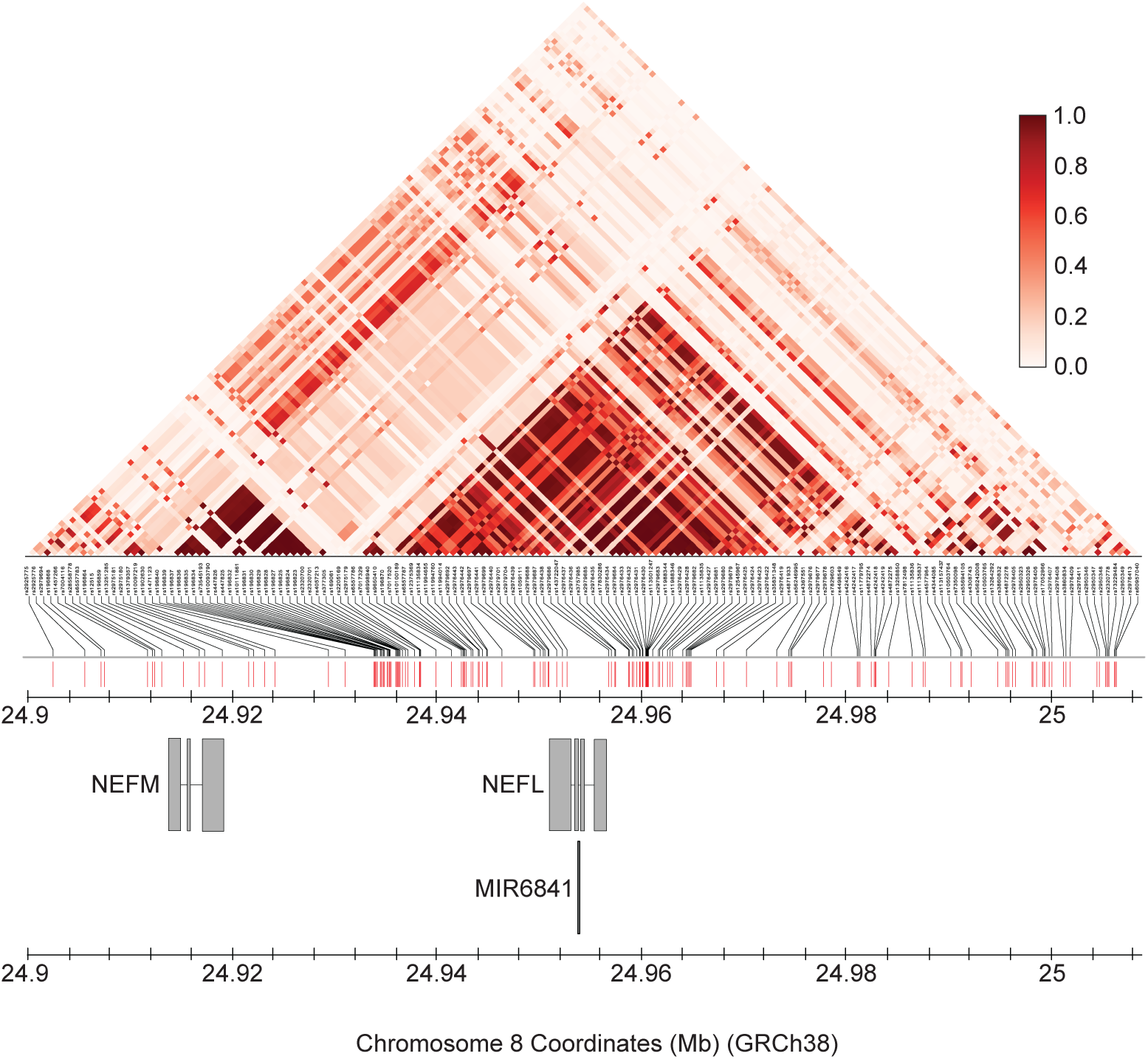
**Linkage measured as R^2 for *NEFL* and 50 kb of flanking DNA on each side**. Linkage values are shown for all SNPs with allele frequency between 0.1 and 0.9 in the region and their position relative to genes in the region is shown below. R^2 values were downloaded from LDLink.

**Figure S19:**
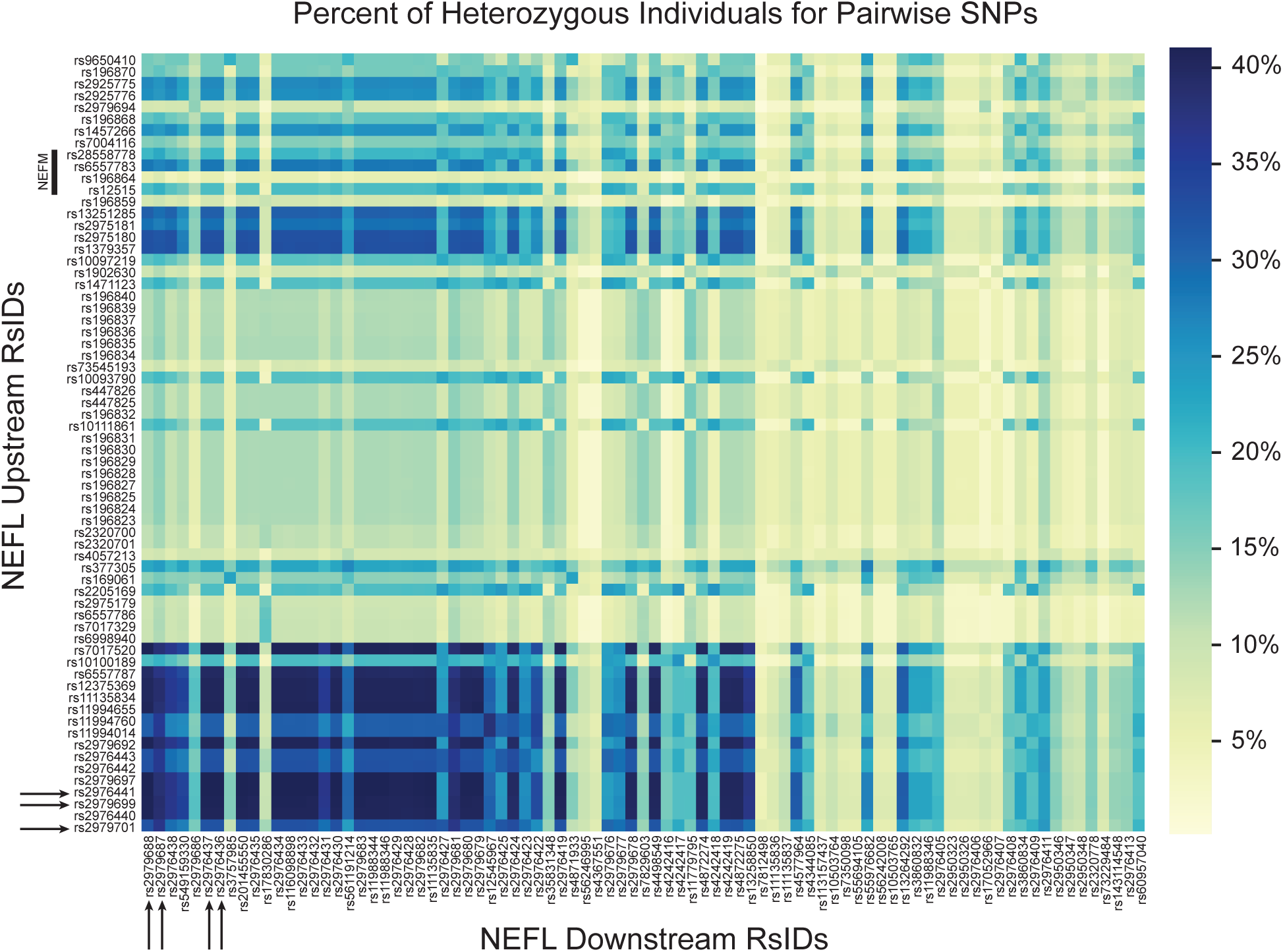
Percentage of individuals heterozygous for pairwise SNPs flanking *NEFL.* Variant data from 1000 Genomes phase 3, starting with minor allele frequency 0.1-0.9 in global population. Horizontal axis indicates SNPs downstream (3’) of *NEFL* increasing in linear distance from left to right. Vertical axis indicates SNPs upstream (5’) of *NEFL* increasing in linear distance from bottom to top. The location of *NEFM* (upstream of *NEFL*) is annotated with a line along the vertical axis. Darker blue colors indicate a higher percentage of heterozygous individuals. Arrows indicate the SNPs targeted in this study.

