## Supplemental Tables for "Haplotype editing with CRISPR/Cas9 as a therapeutic approach for dominant-negative missense mutations in *NEFL*"

### Table 1: Genomic off-target analysis

**N98S gRNA**

| # | chr | start (hg38) | End (hg38) | Strand | CFD score | Description | Results: NF-P3 FS | Results: NF-P3 HDR |
|---|---|---|---|---|---|---|---|---|
| 1 | 1 | 148316436 | 148316458 | + | 0.7143 | intergenic: RP11-495P10.8/RP11-495P10.5/RP11-495P10.7-RP11-495P10.3 | Sequencing Failed | Sequencing Failed |
| 2 | 12 | 5752303 | 5752303 | - | 0.4412 | intron: ANO2 | PCR failed | PCR failed |
| 3 | 10 | 75661072 | 75661094 | - | 0.3956 | intergenic: RP11-310J24.3-RP11-367B6.2 | No mutations | No mutations |
| 4 | 20 | 48137627 | 48137649 | + | 0.3652 | intergenic: AL139351.1-LINC00494 | No mutations | No mutations |
| 5 | 18 | 39020019 | 39020041 | - | 0.3649 | intergenic: RN7SKP182-RNU6-706P | No mutations | No mutations |
| 6 | 4 | 3605528 | 3605550 | - | 0.3563 | intergenic: LINC00955-RP3-368B9.2 | No mutations | No mutations |
| 7 | 9 | 65426450 | 65426472 | + | 0.3563 | intergenic: FOXD4L5-Y_RNA | No mutations | No mutations |
| 8 | 8 | 1925519 | 1925541 | - | 0.3545 | intron: ARHGEF10 | No mutations | No mutations |
| 9 | 13 | 111494882 | 111494904 | + | 0.3254 | intergenic: TEX29-RP11-65D24.2 | No mutations | No mutations |
| 10 | 8 | 120892179 | 120892201 | - | 0.3196 | intergenic: RP11-713M15.2/SNTB1-RP11-369K17.1 | No mutations | No mutations |
| 11 | 9 | 14223924 | 14223946 | + | 0.2790 | intron: NFIB | No mutations | No mutations |

L1.1G and R1.1G

| # | gRNA | chr | start (hg38) | End (hg38) | Strand | CFD score | Description | Results: NF-P3 ex | Results: NF-P3 inv |
|---|---|---|---|---|---|---|---|---|---|
| 1 | R1.1G | 10 | 78632335 | 78632357 | - | 0.49777778 | exon:KCNMA1 | No mutations | No mutations |
| 2 | R1.1G | 3 | 40733428 | 40733450 | + | 0.3799606 | intergenic:RP11-528N21.1-HMGN2P24 | No mutations | No mutations |
| 3 | R1.1G | 5 | 8208980 | 8209002 | + | 0.34821429 | intergenic:RP11-215I16.1-RP11-480D4.1 | No mutations | No mutations |
| 4 | R1.1G | 6 | 18441294 | 18441316 | + | 0.31533434 | intron:RNF144B | No mutations | No mutations |
| 5 | R1.1G | 8 | 48092200 | 48092222 | + | 0.26848498 | intergenic:NDUFA5P12-RP11-1134I14.4 | No mutations | No mutations |
| 6 | R1.1G | 11 | 79198180 | 79198202 | - | 0.38787879 | intergenic:TENM4-RP11-258O13.1 | No mutations | No mutations |
| 7 | R1.1G | 12 | 34271299 | 34271321 | + | 0.38956522 | intergenic:RP11-847H18.3-RP11-313F23.3 | Cell line has a 5 bp variant. No other mutations detected. | Cell line has a 5 bp variant. No other mutations detected. |
| 8 | R1.1G | 4 | 72155681 | 72155703 | + | 0.38181818 | intergenic:SLC4A4-RP11-1J11.1 | No mutations | No mutations |
| 9 | R1.1G | 5 | 133553321 | 133553343 | + | 0.36363636 | intergenic:CTD-2410N18.4/CDKL3/PPP2CA-PPP2CA | No mutations | No mutations |
| 10 | R1.1G | 6 | 68788813 | 68788835 | - | 0.32223776 | intergenic:RP11-542F9.1-RP11-406O16.1 | No mutations | No mutations |
| 11 | R1.1G | 4 | 189766672 | 189766694 | - | 0.30198176 | intergenic:RP11-756P10.2-RP11-818C3.1 | No mutations | No mutations |

L1.1T and R1.1C

| # | gRNA | chr | start (hg38) | End (hg38) | Strand | CFD score | Description | Results: NF-P8 ex | Results: NF-P8 inv |
|---|---|---|---|---|---|---|---|---|---|
| 1 | L1.1T | 5 | 2501128 | 2501150 | + | 0.4982699 | intergenic:Y_RNA- | No mutations | No mutations |
| 2 | L1.1T | 11 | 43929675 | 43929697 | - | 0.29658922 | intergenic:ALKBH3-ALKBH3-AS1 | No mutations | No mutations |
| 3 | L1.1T | 13 | 102135513 | 102135535 | + | 0.27731092 | intron:ITGBL1 | No mutations | No mutations |
| 4 | R1.1C | 12 | 57160521 | 57160543 | + | 0.46928467 | intron:HSD17B6 | No mutations | No mutations |
| 5 | R1.1C | 10 | 92818471 | 92818493 | - | 0.45389474 | intergenic:DDX18P6-LINC00502 | No mutations | No mutations |
| 6 | R1.1C | 3 | 156692992 | 156693014 | - | 0.35 | intergenic:LEKR1-KLF3P2 | No mutations | No mutations |
| 7 | R1.1C | 3 | 80744740 | 80744762 | - | 0.27575758 | intergenic:RP11-481N16.1-RP11-47P18.1 | PCR failed | PCR failed |
| 8 | R1.1C | 3 | 4047631 | 4047653 | + | 0.44426162 | intergenic: PNPT1P1-SUMF1 | No mutations | No mutations |
| 9 | R1.1C | 2 | 57033442 | 57033464 | + | 0.56384439 | intergenic:AC008173.1-snoU13 | Homozygous alternate SNP. No other mutations detected. | Homozygous alternate SNP. No other mutations detected. |

**Table 2: gRNA sequences**

| sgRNA Name | Target | Sequence | Allele Specific (AS) or Biallelic (BA) |
|---|---|---|---|
| N98S | N98S | agctggcgaagcggtcactg | AS |
| E396K | E396K | caggaaacUcUUggaaggcA | AS |
| L1.1G | rs2979688 | GATCACGGCACGCCGGCCAG | AS |
| L1.2G | rs2979688 | AGCGCGCTGCCCCCACTGGC | AS |
| L2T | rs2979687 | TCACGGGGTCTGGGCAATGC | AS |
| L3C | rs2976437 | TCTACATATGGGTAATTGGG | AS |
| L4A | rs2976436 | ACCCATATGTAGATGAAGCA | AS |
| R1.1G | rs2979701 | TCTGTGATAGGTTAGTGTAG | AS |
| R1.2G | rs2979701 | CTGTGATAGGTTAGTGTAGA | AS |
| R2.1T | rs2979699 | TACCAGGGTGACTGGAGTGC | AS |
| R2.2T | rs2979699 | TTCCAGCACTCCAGTCACCC | AS |
| R3G | rs2976441 | GCTTAAATGTCATTCTCTAA | AS |
| L1.1T | rs2979688 | GATCACGGCACGCCGTCCAG | AS |
| L1.2T | rs2979688 | AGCGCGCTGCCCCCACTGGA | AS |
| R1.1C | rs2979701 | TCTGTGATAGGTTAGTCTAG | AS |
| R1.2C | rs2979701 | CTGTGATAGGTTAGTCTAGA | AS |
| BA1 | Intron 1 | gcagctttaatgcggaacgc | BA |
| BA2 | Intron 1 | CCTTTATTTAGTAGGTAGAC | BA |

**Table 3: ssODN sequences for excisions**

| | | |
|---|---|---|
| **L1.1G** | **BA** | CTTGGCTGCAGCAGCGCGCTGCCCCCACTGTTCCGCATTAAAGCTGCCCAGCCCTTGTTG |
| **L1.2G** | **HW15** | CCTTGGCTGCAGCAGCGCGCTGCCCCCACTTTCCGCATTAAAGCTGCCCAGCCCTTGTTG |
| **L1.2G** | **R1.1G** | CCTTGGCTGCAGCAGCGCGCTGCCCCCACTCACTAACCTATCACAGAGTTATAGTGAGAA |
| **L1.1G** | **R1.1G** | CCTTGGCTGCAGCAGCGCGCTGCCCCCACTCACTAACCTATCACAGAGTTATAGTGAGAA |
| **BA** | **R1.1G** | GGGAGTGTGCTCCGTGCTGCTGCACCGGCGCACTAACCTATCACAGAGTTATAGTGAGAA |
| **L1.1T** | **BA** | CTTGGCTGCAGCAGCGCGCTGCCCCCACTGTTCCGCATTAAAGCTGCCCAGCCCTTGTTG |
| **L1.2T** | **BA** | CCTTGGCTGCAGCAGCGCGCTGCCCCCACTTTCCGCATTAAAGCTGCCCAGCCCTTGTTG |
| **L1.2T** | **R1.1C** | CCTTGGCTGCAGCAGCGCGCTGCCCCCACTGACTAACCTATCACAGAGTTATAGTGAGAA |
| **L1.1T** | **R1.1C** | CCTTGGCTGCAGCAGCGCGCTGCCCCCACTGACTAACCTATCACAGAGTTATAGTGAGAA |
| **BA** | **R1.1C** | GGGAGTGTGCTCCGTGCTGCTGCACCGGCGGACTAACCTATCACAGAGTTATAGTGAGAA |

**Table 4: ddPCR assays for excision and inversion frequency**

(exc / inv)

| | R1.1 | R1.2 | R2.1 | R2.2 | R3 |
|---|---|---|---|---|---|
| **L4** | 2 / B | 2 / B | 3 / C | 3 / C | 4 / D |
| **L3** | 2 / B | 2 / B | 3 / C | 3 / C | 4 / D |
| **L2** | 6 / F | 14 / N | 12 / L | 12 / L | 10 / J |
| **L1.2** | 7 / G | 13 / M | 11 / K | 11 / K | 9 / I |
| **L1.1** | 8 / G | 13 / M | 11 / K | 11 / K | 9 / I |

Excision Assays

| **Assay Number** | **F primer** | **R primer** | **Probe** |
|---|---|---|---|
| 2 | HW_ddXR_79_F | HW_ddXR_63_R | HW79 |
| 3 | HW_ddXR_79_F | HW_ddXR_55_R | HW79 |
| 4 | HW_ddXR_79_F | HW_ddXR_53_R | HW79 |
| 6 | HW_ddXR_74_F | HW_ddXR_63_R | HW63 |
| 7 | HW_ddXR_69_F | HW_ddXR_63_R | HW63 |
| 8 | HW_ddXR_72_F | HW_ddXR_63_R | HW63 |
| 9 | CM_ddXR_4790_F | CM_ddXR_11390_R | HW69 |
| 10 | CM_ddXR_4590_F | CM_ddXR_11390_R | HW74 |
| 11 | CM_ddXR_4790_F | CM_ddXR_11270_R | HW69 |
| 12 | CM_ddXR_4590_F | CM_ddXR_11270_R | HW74 |
| 13 | CM_ddXR_4890_F | CM_ddXR_10970_R | HW63 |
| 14 | CM_ddXR_4650_F | CM_ddXR_10970_R | HW63 |

Inversion Assays

| **Assay Letter** | **F primer** | **R primer** | **Probe** |
|---|---|---|---|
| B | HW_ddXR_79_F | CM_iddXR_63_R | HW79 |
| C | HW_ddXR_79_F | CM_iddXR_55_R | HW79 |
| D | HW_ddXR_79_F | CM_iddXR_53_R | HW79 |
| F | CM_iddXR_74_F | HW_ddXR_63_R | HW63 |
| G | CM_NEFL_4996_R | HW_ddXR_63_R | HW63 |
| I | CM_iddXR_11250_F | CM_iddXR_11430_R | HW13 |
| J | CM_ddXR_4590_F | CM_iddXR_4840_R | HW74 |
| K | CM_iddXR_11110_F | CM_iddXR_11310_R | HW13 |
| L | CM_ddXR_4590_F | CM_iddXR_4800_R | HW74 |
| M | CM_iddXR_10800_F | CM_iddXR_10950_R | HW63 |
| N | CM_iddXR_10810_F | CM_iddXR_10950_R | HW63 |

**Table 5: Primer and Probe Sequneces for ddPCR assays for excision and inversion frequency**

Primer Sequences

| Name | Sequence (5' → 3') |
|---|---|
| CM_ddXR_4790_F | ggcatgggatctcagagaaa |
| CM_ddXR_11390_R | caaagaatttgacccactagaagag |
| CM_ddXR_4590_F | gaggttcctggtgggaaa |
| CM_ddXR_11270_R | aaagatgagtgctccagaaa |
| CM_ddXR_4890_F | cgcagaatcctcgcctt |
| CM_ddXR_10970_R | ccctgggagaagggttaga |
| CM_ddXR_4650_F | gaggtgacgggatacagaaa |
| HW_ddXR_79_F | tatgcagactcacacacactg |
| HW_ddXR_72_F | GCAGAATCCTCGCCTTGG |
| HW_ddXR_74_F | GGGCAACTTAAGGATCCAAGT |
| HW_ddXR_63_R | gtggtggcagtataaattgaaaga |
| HW_ddXR_55_R | ATCCTGTGACAGATGGGAGAA |
| HW_ddXR_53_R | TTCAAAGAATTTGACCCACTAGAAG |
| CM_iddXR_74_F | GCTCAGAGGGCCCTGATTTT |
| CM_iddXR_63_R | CTGTGGTCAGTGCCCCTTTT |
| CM_iddXR_55_R | CTGCCTAGTGCTGACTCCTG |
| CM_iddXR_53_R | CTCCACTTCCAGCACTCCAG |
| CM_NEFL_4996_R | ccgttctgccacccctattt |
| CM_iddXR_11250_F | gaggatggatggctgtgtg |
| CM_iddXR_11110_F | gaggatggatggctgtgtg |
| CM_iddXR_10800_F | tatttatacgccgggaggct |
| CM_iddXR_10810_F | ccctcactcatttcccttctg |
| CM_iddXR_11430_R | agatgctaatggcaagaatcaa |
| CM_iddXR_4840_R | ctcccatctgtcacaggattt |
| CM_iddXR_11310_R | aatctgaagggtcagtaggaac |
| CM_iddXR_4800_R | ctgactcctgcctagttctcta |
| CM_iddXR_10950_R | gtggtggcagtataaattgaaagat |

Probe Sequences

| Name | Sequence (5' → 3') |
|---|---|
| HW79 | TGAGGTTTGCAGGGAGCAGGTTAA |
| HW63 | ACACCTCCATGTCTTAGATCCTTCCACA |
| HW13 | caggctgcgtcagg |
| HW69 | tctgagcaaagtggaaaggacgacc |
| HW74 | ctgcgaggtgacgggatacagaaa |

**Table 6: Antibodies**

| **Name/Antigen** | **Host** | **Vendor/Catalog #** | **IF Concentration** |
|---|---|---|---|
| NF-L | Rabbit | millipore<br>AB9568 | 1:1000 for CX7,<br>1:500 for Keyence |
| β-Tubulin III | Mouse | ThermoFisher<br>480011 | 1:250 |
| HB9 | Mouse | DSHB 81.5c10 | 1:200 |
| Anti-rabbit<br>Alexa Fluor 488 | Goat | Invitrogen A11034 | 1:500 |
| Anti-mouse<br>Alexa Fluor 594 | Goat | Invitrogen A11032 | 1:500 |

**Table 7 : Assays for inversions and excisions**

| Excision Frequency | | | |
|---|---|---|---|
| Guide Pair | F Primer | R Primer | Probe |
| L1.1 - BA | PD_ddXR_72/73F_anchored | PD_ddXR_HW15_R | HW63 |
| L1.1 - R1.1 | HW_ddXR_72_F | HW_ddXR_63_R | HW63 |
| BA - R1.1 | HW_ddXR_15/63F | HW_ddXR_63_R | HW63 |

| Inversion Frequency | | | |
|---|---|---|---|
| Guide Pair | F Primer | R Primer | Probe |
| L1.1 - BA | PD_ddXR_72/73F_anchored | PD_iddXR_HW15_R | HW63 |
| L1.1 - R1.1 | CM_NEFL_4996_R | HW_ddXR_63_R | HW63 |
| BA - R1.1 | HW_iddXR_15/63F | HW_ddXR_63_R | HW63 |

| Excision Specificity via ddPCR | | | |
|---|---|---|---|
| Guide Pair | F Primer | R Primer | Allele Discrimination Assay |
| L1.1 - BA | NEFL_forASaroundrs2979685 | PD_7984_R | rs2979685 |
| L1.1 - R1.1 | NEFL_forASaroundrs2979685 | HW_ddXR_63_R | rs2979685 |
| BA - R1.1 | NEFL_forASaroundrs2979685 | HW_ddXR_63_R | rs2979685 |

| Inversion Specificity via ddPCR | | | |
|---|---|---|---|
| Guide Pair | F Primer | R Primer | Allele Discrimination Assay |
| L1.1 - BA | NEFL_forASaroundrs2979685 | HW_NEFL_5740_F | rs2979685 |
| L1.1 - R1.1 | NEFL_forASaroundrs2979685 | BMJS_NEFL_7758F | rs2979685 |
| BA - R1.1 | NEFL_forASaroundrs2979685 | HW_ddXR_63_R | rs2979685 |

**Table 8: Primer sequences for excision and inversion assays**

| Name | Sequence 5' → 3' |
|---|---|
| PD_ddXR_72/73F_anchored | CCCTCTGAGCAAAGTGGAAA |
| HW_ddXR_15/63F | TGTAGTCTGGGAGTGTGCT |
| HW_iddXR_15/63F | CCCAATTCCCACGTCTTCC |
| NEFL_forASaroundrs2979685 | GGCTGTCGTGGAGTATGAGG |
| PD_ddXR_HW15_R | CAGGTCAGTAGAGAGCTGAT |
| HW_ddXR_63_R | GTGGTGGCAGTATAAATTGAAAGA |
| PD_iddXR_HW15_R | TGTAGTCTGGGAGTGTGCT |
| PD_7984_R | CTCCTCTTGGACATGGCTGG |
| HW_ddXR_72_F | GCAGAATCCTCGCCTTGG |
| CM_NEFL_4996_R | CCGTTCTGCCACCCCTATTT |
| HW_NEFL_5740F | TCGACAGCTTGATGGACGAA |
| BMJS_NEFL_7758F | TCCTGCTTGCCTTTGTGTTTAG |

**Table 9 : hNIL copy number 20x primer probe mixes**

| hNIL copy number ddPCR custom mixes | | | |
|---|---|---|---|
| | F Primer | R Primer | Probe |
| Neomycin Assay | catggctgatgcaatgcg | tcgcttggtggtcgaatg | cgcttgatccggctacctgcc |
| TRE3G Assay | tacggtgggcgcctataa | agtggtacggaaagttggtataag | agatcgcctggagcaattccacaa |

**Table 10: hNIL genotyping junction PCR primer sequences**

| Junction PCR Primers for hNIL CLYBL integration | |
|---|---|
| hNIL CLYBL 5’ Junction F | CAGACAAGTCAGTAGGGCCA |
| hNIL CLYBL 5’ Junction R | AGAAGACTTCCTCTGCCCTC |
| hNIL CLYBL 3’ Junction F | CACCAGCAACCTGACGTTTT |
| hNIL CLYBL 3’ Junction R | TTTTATAGGCGCCCACCGTA |
| CLYBL WT F | TGACTAAACACTGTGCCCCA |
| CLYBL WT R | AGGCAGGATGAATTGGTGGA |

| Post-Cre Junction PCR Primers | |
|---|---|
| CF28 | CCTCAGCCCAGTTTCCACTTG |
| CF29 | GGCTATGAACTAATGACCCCGT |

**Table 11 : Primer sequences for next generation sequencing**

| Name | Sequence 5' → 3' with illumina adapter sequence |
|---|---|
| BMJS_NGS_10733F | TCCCTACACGACGCTCTTCCGATCTcagctAgttctctttcagtattccctctcc |
| BMJS_NGS_10965R | GTTCAGACGTGTGCTCTTCCGATCTcagctAggttagagtggtggcagtataaa |
| BMJS_NGS_4810F | TCCCTACACGACGCTCTTCCGATCTcagctAccctctgagcaaagtggaaa |
| BMJS_NGS_5025R | GTTCAGACGTGTGCTCTTCCGATCTcagctAgaggatggatggctgtgtg |
